# Uniparental nuclear inheritance following bisexual mating in fungi

**DOI:** 10.1101/2020.12.03.410415

**Authors:** Vikas Yadav, Sheng Sun, Joseph Heitman

## Abstract

Some remarkable animal species require an opposite-sex partner for their sexual development but discard the partner’s genome before gamete formation, generating hemi-clonal progeny in a process called hybridogenesis. Here, we discovered a similar phenomenon, termed pseudosexual reproduction, in a basidiomycete human fungal pathogen, *Cryptococcus neoformans*, where exclusive uniparental inheritance of nuclear genetic material was observed during bisexual reproduction. Analysis of strains expressing fluorescent reporter proteins revealed instances where only one of the parental nuclei was present in the terminal sporulating basidium. Whole-genome sequencing revealed the nuclear genome of the progeny was identical with one or the other parental genome. Pseudosexual reproduction was also detected in natural isolate crosses where it resulted in mainly *MAT*α progeny, a bias observed in *Cryptococcus* ecological distribution as well. The meiotic recombinase Dmc1 was found to be critical for pseudosexual reproduction. These findings reveal a novel, and potentially ecologically significant, mode of eukaryotic microbial reproduction that shares features with hybridogenesis in animals.

## Introduction

Most multicellular organisms in nature undergo (bi)sexual reproduction involving two partners of the opposite sex to produce progeny. In most cases, following fusion of the two haploid gametes, the diploid zygote receives one copy of the genetic material from each parent. To produce these haploid gametes, a diploid germ cell of the organism undergoes meiosis, which involves recombination between the two parental genomes, generating recombinant progeny. Recombination confers benefits by bringing together beneficial mutations and segregating away deleterious ones (Dimijian, 2005; Meirmans, 2009). In contrast, some organisms undergo variant forms of sexual reproduction, including parthenogenesis, gynogenesis, androgenesis, and hybridogenesis, and, in doing so, produce clonal or hemi-clonal progeny (Avise, 2015; Neaves & Baumann, 2011).

In parthenogenesis, a female produces clonal progeny from its eggs without any contribution from a male partner (Avise, 2015; Hörandl, 2009). Gynogenesis and androgenesis occur when the fusion of an egg with a sperm induces cell division to produce clonal female or male zygotes, respectively (Lehtonen, Schmidt, Heubel, & Kokko, 2013). During hybridogenesis, an egg from one species fuses with the sperm from another species to generate a hybrid diploid zygote (Lavanchy & Schwander, 2019). However, one of the parental genomes is excluded during development, in a process termed genome exclusion that occurs before gametogenesis. The remaining parental genome undergoes replication followed by meiosis to produce an egg or a sperm. The sperm or egg then fuses with an opposite-sex gamete to generate a hemiclonal progeny. Because only one parent contributes genetic material to the progeny, but both parents are physically required, this phenomenon has been termed sexual parasitism (Lehtonen et al., 2013; Umphrey, 2006). While most of the reported cases of hybridogenesis are from female populations, recent reports suggest that it may also occur in male populations of some species (Dolezalkova et al., 2016; Schwander & Oldroyd, 2016). Currently hybridogenesis has only been observed in the animal kingdom in some species of frogs, fishes, and snakes. Plants also exhibit parthenogenesis (aka apomixis), along with gynogenesis and androgenesis (Lehtonen et al., 2013; Mirzaghaderi & Horandl, 2016).

Unlike animals, most fungi do not have sex chromosomes; instead, cell-type identity is defined by the mating-type (*MAT*) locus (Heitman, 2015; Heitman, Sun, & James, 2013). While many fungi are heterothallic, with opposite mating-types in different individuals, and undergo sexual reproduction involving two partners of compatible mating-types, other fungi are homothallic, with opposite mating-types within the same organism, and can undergo sexual production during solo culture in the absence of a mating partner. One class of homothallic fungi undergoes unisexual reproduction, during which cells of a single mating type undergo sexual reproduction to produce clonal progeny, similar to parthenogenesis (Heitman, 2015; Lee, Ni, Li, Shertz, & Heitman, 2010). Gynogenesis and hybridogenesis have not been identified in the fungal kingdom thus far.

*Cryptococcus neoformans* is a basidiomycete human fungal pathogen that exists as either one of two mating types, *MAT***a** or *MAT*α (Sun, Coelho, David-Palma, Priest, & Heitman, 2019). During sexual reproduction, two haploid yeast cells of opposite mating type interact and undergo cell-cell fusion (Kwon-Chung, 1975, 1976; Sun, Priest, & Heitman, 2019). The resulting dikaryotic zygote then undergoes a morphological transition and develops into hyphae whose termini mature to form basidia. In the basidium, the two parental nuclei fuse (karyogamy), and the resulting diploid nucleus undergoes meiosis to produce four daughter nuclei (Idnurm, 2010; Kwon-Chung, 1976; Sun, Priest, et al., 2019; Zhao, Lin, Fan, & Lin, 2019). These four haploid nuclei repeatedly divide via mitosis and bud from the surface of the basidium to produce four long spore chains. Interestingly, apart from this canonical heterothallic sexual reproduction, a closely related species, *C. deneoformans* can undergo unisexual reproduction (Lin, Hull, & Heitman, 2005; Roth, Sun, Billmyre, Heitman, & Magwene, 2018; Sun, Billmyre, Mieczkowski, & Heitman, 2014).

In a previous study, we generated a genome-shuffled strain of *C. neoformans*, VYD135α, by using the CRISPR-Cas9 system targeting centromeric transposons in the lab strain H99α. This led to multiple centromere-mediated chromosome arm exchanges in strain VYD135α when compared to the parental strain H99α, without any detectable changes in gene content in the two genomes (Yadav, Sun, Coelho, & Heitman, 2020). Additionally, strain VYD135α exhibits severe sporulation defects when mated with strain KN99**a** (which is congenic with strain H99α but has the opposite mating type) likely due to the extensive chromosomal rearrangements introduced into the VYD135α strain. In this study, we show that the genome-shuffle strain VYD135α can in fact produce spores in crosses with *MAT***a** *C. neoformans* strains after prolonged mating. Analysis of these spores reveals that the products from an individual basidium contain genetic material derived from only one of the two parents. Whole-genome sequencing of the progeny revealed an absence of recombination between the two parental genomes. The mitochondria in these progeny were found to always be inherited from the *MAT***a** parent, consistent with known mitochondrial uniparental inheritance (UPI) patterns in *C. neoformans* (Sun, Fu, Ianiri, & Heitman, 2020). Using strains with differentially fluorescently labeled nuclei, we discovered that in a few hyphal branches as well as in basidia, only one of the two parental nuclei was present and produced spores, termed uniparental nuclear inheritance. We also observed the occurrence of such uniparental nuclear inheritance in wild-type and natural isolate crosses. Furthermore, we found that the meiotic recombinase Dmc1 plays a central role during this unusual mode of reproduction of *C. neoformans*. Overall, this mode of sexual reproduction of *C. neoformans* exhibits striking parallels with hybridogenesis in animals.

## Results

### Chromosomal translocation strain exhibits unusual sexual reproduction

Previously, we generated a strain (VYD135α) with eight centromere-mediated chromosome translocations compared to the wild-type parental isolate H99α (Yadav et al., 2020). Co-incubation of the wild-type strain KN99**a** with the genome-shuffle strain VYD135α resulted in hyphal development and basidia production, but no spores were observed during a standard two-week incubation. However, when sporulation was assessed at later time points in the VYD135α × KN99**a** cross, we observed a limited number of sporulating basidia (16/1201=1.3%) after five weeks of mating compared to much greater level sporulation in the wild-type H99α × KN99**a** cross (524/599 = 88%) (Figure 1A–D). None of these strains exhibited any filamentation on their own even after 5-weeks of incubation, indicating the sporulation events are not a result of unisexual reproduction (Figure 1A–B). To analyze this delayed sporulation process in detail, spores from individual basidia were dissected and germinated to yield viable F1 progeny. As expected, genotyping of the mating-type locus in the H99α × KN99**a** progeny revealed the presence of both mating types in spores derived from each basidium (Figure 1E and G, Table 1). On the other hand, the same analysis for VYD135α × KN99**a** revealed that all germinating progeny from each individual basidia possessed either only the *MAT*α or the *MAT***a** alleles (Figure 1E and G, Table 1). PCR assays also revealed that the mitochondria in all of these progeny were inherited from the *MAT***a** parent, in accord with known UPI (Figure 1F–G). These results suggest the inheritance of only one of the parental nuclei in the VYD135α × KN99**a** F1 progeny. The presence of mitochondria from the *MAT***a** parent further confirmed that these progeny were not the products of unisexual reproduction.

**Table 1.** Genotype analysis of basidia-specific spores germinated from H99α × KN99a and VYD135α × KN99a crosses.

| Basidia<br># | H99α x KN99a cross | | | | VYD135α x KN99a cross | | | |
| --- | --- | --- | --- | --- | --- | --- | --- | --- |
|  | Spores<br>germinated/<br>dissected | %<br>germinated | *MAT* | Mito | Spores<br>germinated/<br>dissected | %<br>germinated | *MAT* | Mito |
| 1 | 5/14 | 36 | 4α + 1a | a | 12/24 | 50 | All $\alpha$ | a |
| 2 | 14/14 | 100 | 7α + 7a | a | 6/10 | 60 | All $\alpha$ | a |
| 3 | 12/14 | 86 | 2α + 7a<br>+ 3a/$\alpha$ | a | 15/15 | 100 | All a | a |
| 4 | 10/14 | 71 | 4α + 6a | a | 22/27 | 81 | All a | a |
| 5 | 7/13 | 54 | 6a + 1a/$\alpha$ | a | 3/12 | 25 | All $\alpha$ | a |
| 6 | 13/14 | 93 | 6α + 7a | a | 25/27 | 93 | All $\alpha$ | a |
| 7 | 11/14 | 79 | 6α + 5a | a | 4/4 | 100 | All $\alpha$ | a |
| 8 | 14/14 | 100 | 12α + 2a | a | 10/13 | 77 | All $\alpha$ | a |
| 9 | 10/14 | 71 | 4α + 6a | a | 13/15 | 87 | All $\alpha$ | a |
| 10 | 14/14 | 100 | 7α + 7a | a | 31/61 | 51 | All $\alpha$ | a |
| 11 | 14/14 | 100 | 10α + 4a | a | 10/10 | 100 | All a | a |
| 12 | 12/14 | 86 | 8α + 4a | a | 4/5 | 80 | All a | a |
| 13 | 4/11 | 36 | All a | a | 24/28 | 86 | All a | a |
| 14 | 13/13 | 100 | 8α + 5a | a | 16/28 | 57 | All a | a |
| 15 | 14/14 | 100 | 7α + 7a | a | 11/11 | 100 | All a | a |
| 16 | 14/14 | 100 | 6α + 8a | a | 10/22 | 45 | All $\alpha$ | a |

- 3 Mito refers to Mitochondria.

**Figure 1.**
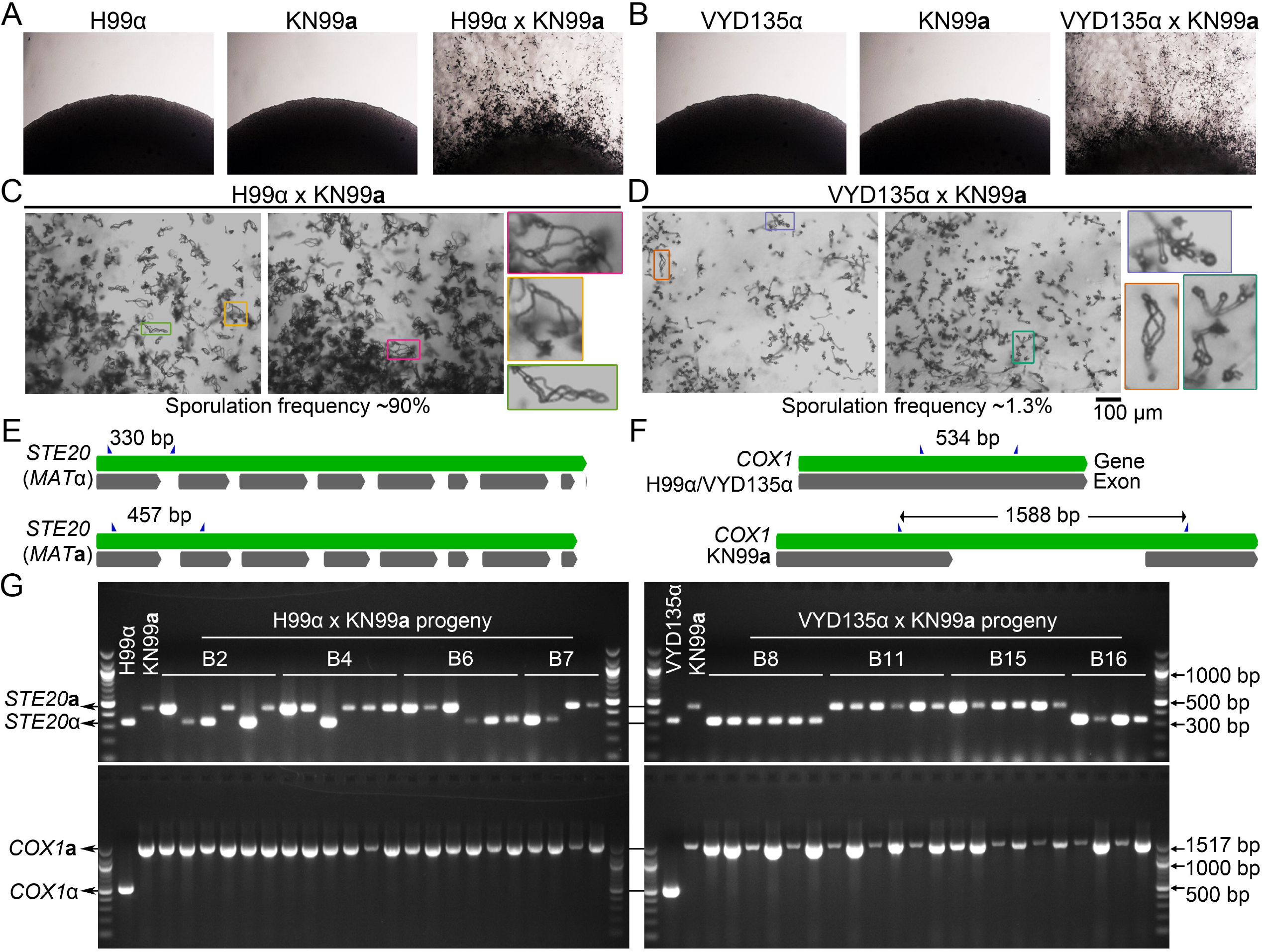
Chromosome shuffled strain exhibits unusual sexual reproduction. **(A-B)** Images of cultures for the individual strains showing no self-filamentation on mating medium. Magnification=10X. **(C-D)** Light microscopy images showing robust sporulation in the H99α × KN99**a** cross, whereas the VYD135α × KN99**a** cross exhibited infrequent sporulation events. The inset images show examples of basidia observed in each of the crosses. Bars, 100 μm. **(E-F)** A scheme showing the *MAT*α (H99α and VYD135α) and *MAT***a** (KN99**a**) alleles at the *STE20* **(E)** and *COX1* **(F)** loci. Primers used for PCR analysis are marked by blue triangles. **(G)** Gel images showing PCR amplification of *STE20* and *COX1* alleles in the progeny obtained from four different basidia for both H99α × KN99**a** and VYD135α × KN99**a** crosses. PCR analysis for the parental strains is also shown and key bands for DNA marker are labeled.

### Fluorescence microscopy reveals uniparental nuclear inheritance after mating

Next, we tested whether the uniparental inheritance detected at the *MAT* locus also applies to the entire nuclear genome. To address this, we established a fluorescence-based assay where the nucleus of strains H99α and VYD135α was labeled with GFP-H4, whereas the KN99**a** nucleus was marked with mCherry-H4. In a wild-type cross (H99α × KN99**a**), the nuclei in the hyphae as well as in the spores are yellow to orange because both nuclei are in a common cytoplasm and thus incorporate both the GFP- and the mCherry-tagged histone H4 proteins (Figure S1A and B). We hypothesized that in the cases of uniparental nuclear inheritance, only one of the nuclei would reach the terminal basidium and thus would harbor only one fluorescent nuclear color signal (Figure S1A).

After establishing this fluorescent tagging system using wild-type strains, H99α × KN99**a** and shuffle-strain VYD135α × KN99**a** crosses with fluorescently labeled strains were examined. In the wild-type cross, most of the basidia formed robust spore chains with both fluorescent colors observed in them while a small population (~1%) of basidia exhibited spore chains with only one color, representing uniparental nuclear inheritance (Figure 2A and S2A). On the other hand, the majority of the basidia population in the shuffle-strain VYD135α × KN99**a** cross did not exhibit sporulation, and the two parental nuclei appeared fused but undivided (Figure 2B and S2B). A few basidia (~1%) bore spore chains with only one fluorescent color, marking uniparental nuclear inheritance events. While the basidia with uniparental nuclear inheritance in the H99α × KN99**a** cross were a small fraction (~1%) of sporulating basidia, the uniparental basidia accounted for all of the sporulating basidia in the VYD135α × KN99**a** cross. Taken together, these results show that the uniparental nuclear inheritance leads to the generation of clonal progeny but requires mating, cell-cell fusion between parents of two opposite mating types. Thus, this process defies the main purpose of sexual reproduction, which is to produce recombinant progeny from two parents. Based on these observations, we define the process of uniparental nuclear inheritance during mating in *C. neoformans* as pseudosexual reproduction (and it is referred to as such hereafter). The progeny obtained via this process will be referred to as the uniparental progeny because they inherit a nuclear genome derived from only one of the two parents.

**Figure 2.**
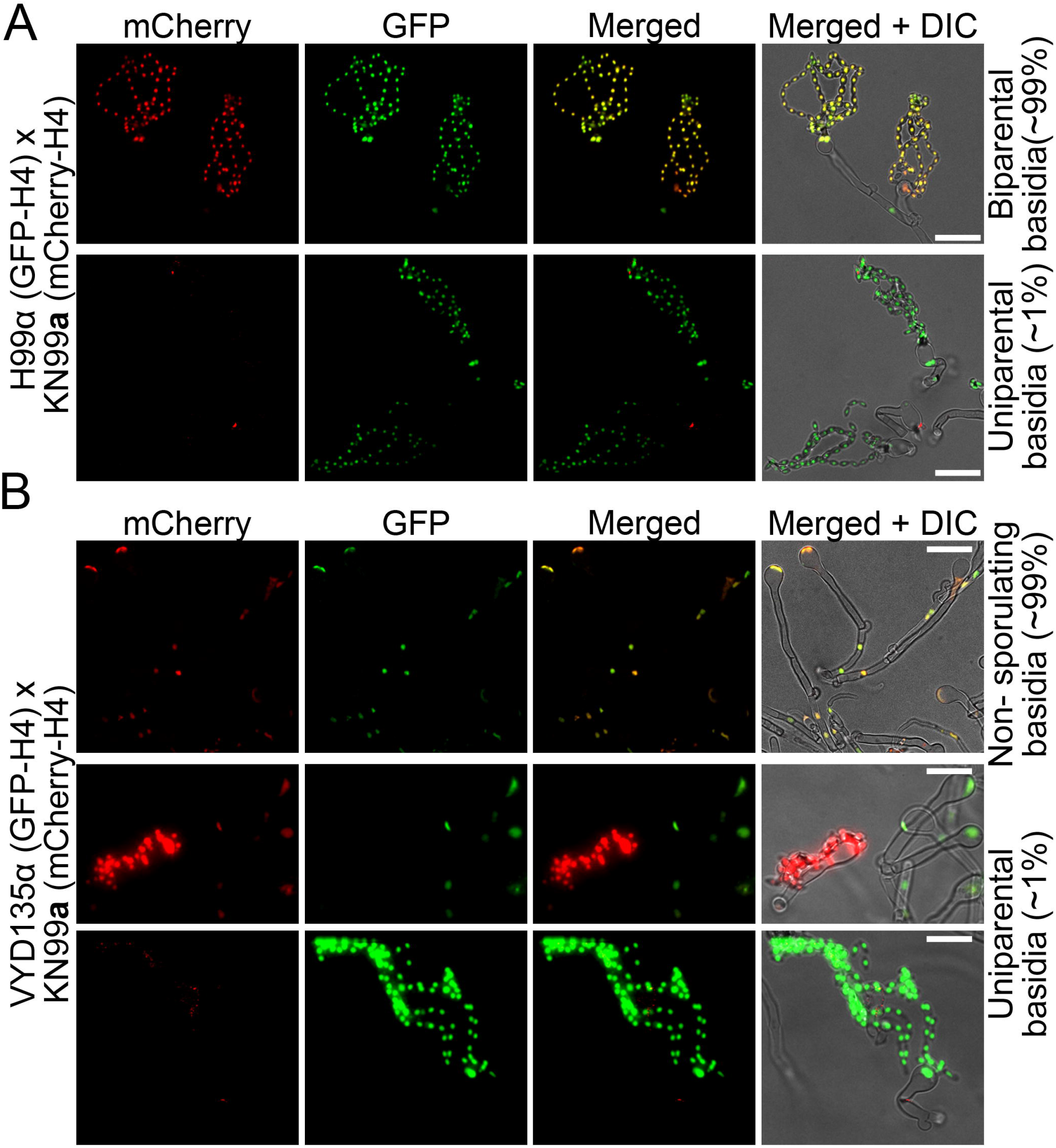
Fluorescence microscopy reveals uniparental nuclear inheritance in the wild-type crosses. **(A)** Mating of GFP-H4 tagged H99α and mCherry-H4 tagged KN99**a** revealed the presence of both fluorescent markers in most spore chains along with uniparental nuclear inheritance in rare cases (~1%). In these few sporulating basidia, only one of the fluorescent signals was observed in the spore chains, reflecting the presence of only one parental nucleus in these basidia. **(B)** Crosses involving GFP-H4 tagged VYD135α, and mCherry-H4 tagged KN99**a** revealed the presence of spore chains with only one fluorescent color. In the majority of basidia that have both parental nuclei, marked by both GFP and mCherry signals, spore chains are not produced suggesting a failure of meiosis in these basidia. Bars, 10 μm.

### Pseudosexual reproduction also occurs in natural isolates

After establishing the pseudosexual reproduction of lab strains, we sought to determine whether such events also occur with natural isolates. For this purpose, we selected two wild-type natural isolates, Bt63**a** and IUM96-2828**a** (referred to as IUM96**a** hereafter) (Desjardins et al., 2017; Keller, Viviani, Esposto, Cogliati, & Wickes, 2003; Litvintseva et al., 2003). IUM96**a** belongs to the same lineage as H99α/KN99**a** (VNI) and exhibits approximately 0.1% genome divergence from the H99α reference genome. Bt63**a** belongs to a different lineage of the *C. neoformans* species (VNBI) and exhibits ~0.5% genetic divergence from the H99α/KN99**a** genome. Both the Bt63**a** and the IUM96**a** genomes exhibit one reciprocal chromosome translocation with H99α and, as a result, share a total of ten chromosome-level changes with the genome-shuffle strain VYD135α (Figure 3A). None of these strains are self-filamentous even after prolonged incubation on mating media but both mate efficiently with both H99α and VYD135α (S3A).

**Figure 3.**
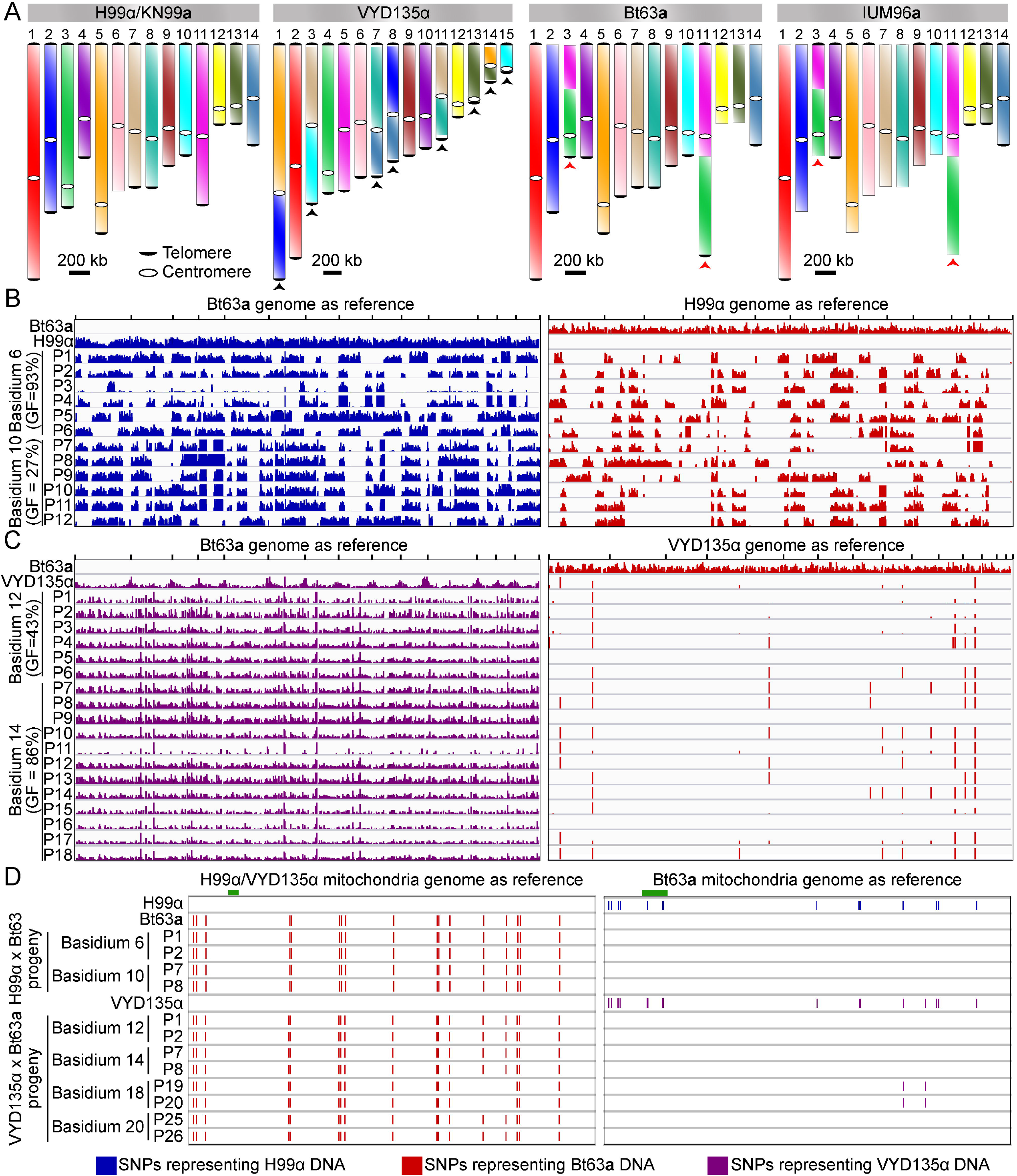
VYD135α progeny exhibit strict uniparental nuclear inheritance and lack the signature of meiotic recombination. **(A)** Chromosome maps for H99α/KN99**a**, VYD135α, Bt63**a**, and IUM96**a** showing the karyotype variation. The genome of the wild-type strain H99α served as the reference. Black arrowheads represent chromosome translocations between VYD135α and H99α whereas red arrowheads mark chromosomes with a translocation between H99α and Bt63**a** or IUM96**a**. **(B)** Whole-genome sequencing, followed by SNP identification, of H99α × Bt63**a** progeny revealed evidence of meiotic recombination in all of the progeny. The left panel shows SNPs with respect to the Bt63**a** genome whereas the right panel depicts SNPs against the H99α genome. H99α and Bt63**a** Illumina sequencing data served as controls for SNP calling. **(C)** SNP analysis of VYD135α × Bt63**a** progeny revealed no contribution of the Bt63**a** parental genome in the progeny as evidenced by the presence of SNPs only against Bt63**a** (left panel) but not against the VYD135α genome (right panel). The presence of a few SNPs observed in VYD135α, as well as all VYD135α × Bt63**a** progeny, are within nucleotide repeat regions. GF stands for germination frequency and P stands for progeny. **(D)** SNP analysis of H99α × Bt63**a** and VYD135α × Bt63**a** progeny using mitochondrial DNA as the reference revealed that mitochondrial DNA is inherited from Bt63**a** in all of the progeny. Progeny obtained from VYD135α × Bt63**a** basidium 18 also revealed recombination between the two parental mitochondrial genomes as marked by the absence or presence of two SNPs when mapped against VYD135α and Bt63 mitochondrial genomes, respectively. The green bar in each panel depicts the locus used for PCR analysis of the mitochondrial genotype in the progeny.

During mating, the H99α × Bt63**a** cross rapidly (within a week) producing robust sporulation from most of the basidia observed. The VYD135α × Bt63**a** cross underwent a low frequency of sporulation (12 spore-producing basidia/840 basidia=1.4%) in 2 to 3 weeks (Figure S3B). Dissection of spores from the H99α × Bt63**a** cross revealed a low germination frequency (average of 25%) with two of the basidia showing no spore germination at all (Table S1). This result is consistent with previous results and the low germination frequency could be explained by the genetic divergence between the two strains (Morrow et al., 2012). Genotyping of germinated spores from the H99α × Bt63**a** cross revealed both *MAT***a** and *MAT*α progeny from individual basidia, with almost 75% of the meiotic events generating progeny that were heterozygous for the *MAT* locus (Figure S3C and Table S1). For the VYD135α × Bt63**a** cross, spores from 15/20 basidia germinated and displayed higher germination frequency than the H99α × Bt63**a** cross (Table S1). Interestingly, all germinated progeny harbored only the *MAT*α mating-type whereas the mitochondria were in all cases inherited from the *MAT***a** parent (Figure S3C). These results suggest pseudosexual reproduction also occurs with Bt63**a** and accounts for the high germination frequency of progeny from the VYD135α × Bt63**a** cross. The occurrence of pseudosexual reproduction was also identified using the fluorescence-based assay with crosses between the GFP-H4 tagged VDY135α and mCherry-H4 tagged Bt63**a** strains (Figure S4).

Mating assays with strain IUM96**a** also revealed a low level of sporulation (19/842=2.3%) with VYD135α but a high sporulation frequency with H99α (91%) (Figure S3D). Analysis of progeny from crosses involving IUM96**a** revealed a similar pattern to what was observed with crosses involving KN99**a**. The progeny from H99α × IUM96**a** exhibited variable basidia-specific germination frequency and inherited both *MAT***a** and *MAT*α in each basidium, whereas VYD135α × IUM96**a** progeny from each basidium inherited exclusively either *MAT***a** or *MAT*α (Figure S3E, Table S2). Interestingly, we observed co-incident uniparental *MAT* inheritance and a high germination frequency in progeny of basidia 7, 8, and 9 from the H99α × IUM96**a** cross as well (Figure S3E, Table S2). Taken together, these results suggest that this unusual mode of sexual reproduction occurs with multiple natural isolates. We further propose that pseudosexual reproduction occurs in nature in parallel with standard sexual reproduction.

### Uniparental progeny completely lack signs of recombination between the two parents

As mentioned previously, H99α (as well as the H99α-derived strain VYD135α) and Bt63**a** have approximately 0.5% genetic divergence. The occurrence of pseudosexual reproduction in the VYD135α × Bt63**a** cross allowed us to test if the two parental genomes recombine with each other during development. We subjected some of the VYD135α × Bt63**a**, as well as the H99α × Bt63**a**, progeny to whole-genome sequencing. As expected, for the H99α × Bt63**a** cross, both parents contributed to the nuclear composition of their progeny, and there was clear evidence of meiotic recombination as determined by variant analysis (Figure 3B). When the VYD135α × Bt63**a** progeny were similarly analyzed, the nuclear genome in each progeny was found to be inherited exclusively from only the VYD135α parent (Figure 3C and S5), and the progeny exhibited sequence differences across the entire Bt63**a** genome. In contrast, the mitochondrial genome was inherited exclusively from the Bt63**a** parent (Figure 3D and S6), in accord with the PCR assay results discussed above. Additionally, the whole-genome sequencing data also revealed that while most of the H99α × Bt63**a** progeny exhibited aneuploidy, the genome-shuffle strain VYD135α × Bt63**a** progeny were euploid (Figure S7A-B), and based on flow cytometry analysis these uniparental progeny were haploid (Figure S7C).

The progeny from crosses involving IUM96**a** as the *MAT***a** partner were also sequenced. Similar to the Bt63**a** analysis, the H99α × IUM96**a** progeny exhibited signs of meiotic recombination, whereas the VYD135α × IUM96**a** progeny did not (Figure S8). Congruent with the mating-type analysis, the progeny in each of the basidia exclusively inherited nuclear genetic material from only one of the two parents. Furthermore, the H99α × IUM96**a** progeny were found to be aneuploid for some chromosomes while the progeny of VYD135α × IUM96**a** were completely euploid (Figure S9). We also sequenced four progeny from basidium 7 from the H99α × IUM96**a** cross, which were suspected to be uniparental progeny based on mating-type PCRs. This analysis showed that all four progeny harbored only H99α nuclear DNA and had no contribution from the IUM96**a** nuclear genome, further supporting the conclusion that pseudosexual reproduction occurs in wild-type crosses (Figure S8A). Similar to other progeny, the mitochondria in these progeny were inherited from the *MAT***a** parent (Figure S3E and Table S2). Combined, these results affirm the occurrence of a novel mode of sexual reproduction in *C. neoformans*, which is initiated by two strains of opposite mating types, but only one of the two parental nuclei is retained during sexual development to eventually form basidia and produce basidiospores through pseudosexual reproduction.

### Pseudosexual reproduction stems from nuclear loss via hyphal branches

Fluorescence microscopy revealed that only one of the two parental nuclei is present in a small proportion of the basidia, which results in meiosis and sporulation. Based on this finding, we hypothesized that the basidia with only one parental nucleus might arise due to nuclear segregation events during hyphal branching. To gain further insight into this process, the nuclear distribution pattern along the sporulating hyphae was studied. As expected, imaging of long hyphae in the wild-type cross revealed the presence of pairs of nuclei with both fluorescent markers along the length of the majority of hyphae (Figure 4A). In contrast, tracking of hyphae from basidia with spore chains in the genome-shuffle strain VYD135α × KN99**a** cross revealed hyphal branches with only one parental nucleus, which were preceded by a hypha with both parental nuclei (Figure 4B, S10A and B). Unfortunately, a majority of the hyphae we tracked were embedded into the agar and most of these could not be tracked to the point of branching. For some others, we were able to image the hyphal branching point where two nuclei separate from each other but were then either broken or did not have mature basidia on them (Figure S10B). We also observed long hyphae with only one parental nucleus in VYD135α × Bt63**a** cross as well, suggesting the mechanism might be similar between strains.

**Figure 4.**
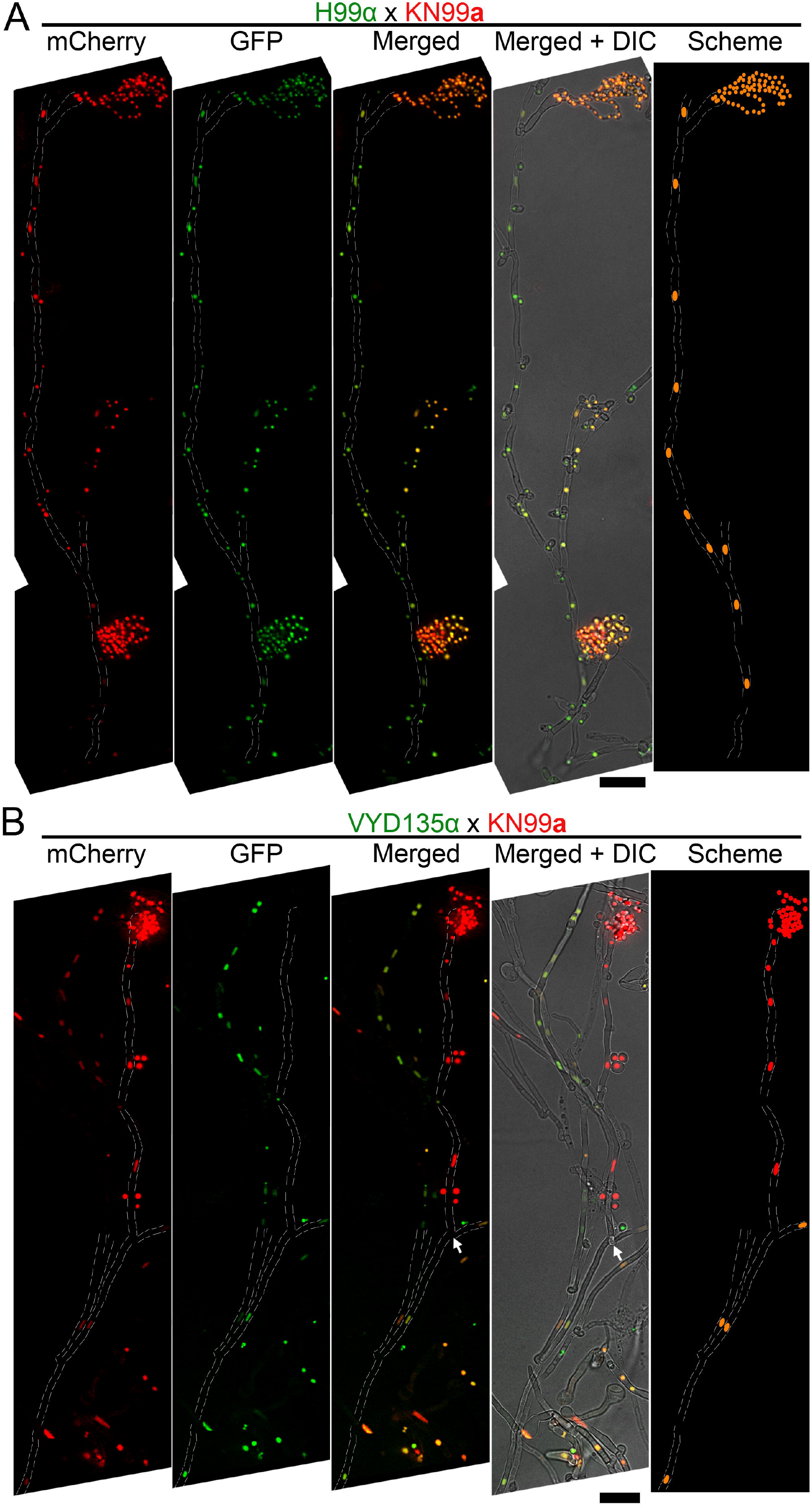
Pan-hyphal microscopy reveals the loss of one parental nucleus during pseudosexual reproduction. Spore-producing long hyphae were visualized in both (**A**) wild-type H99α × KN99**a** and (**B**) VYD135α × KN99**a** crosses to study the dynamics of nuclei in hyphae. Both nuclei were present across the hyphal length in the wild-type and resulted in the production of recombinant spores. On the other hand, one of the nuclei was lost during hyphal branching in the VYD135α × KN99**a** cross and resulted in uniparental nuclear inheritance in the spores that were produced. The arrow in **B** marks the hyphal branching point after which only one of the parental nuclei is present (also see figure S10A). The images were captured as independent sections and assembled to obtain the final presented image. Bars, 10 μm.

These results suggest that hyphal branching may facilitate the separation of one parental nucleus from the main hyphae harboring both parental nuclei. While this is the most plausible explanation based on our results, we cannot rule out other possible mechanisms, such as a role for clamp cells, leading to nuclear separation during hyphal growth. As a result, one of the parental genomes is excluded at a step before diploidization and meiosis, similar to the process of genome exclusion observed in hybridogenesis. We hypothesize that nuclear segregation can be followed by endoreplication occurring in these hyphal branches or in the basidia to produce a diploid nucleus that then ultimately undergoes meiosis and produces uniparental progeny.

### Meiotic recombinase Dmc1 is important for pseudosexual reproduction

Because the genomes of the uniparental progeny did not show evidence of meiotic recombination between the two parents, we sought to test whether pseudosexual reproduction involves meiosis. Additionally, we sought to obtain evidence for our hypothesis that pseudosexual reproduction involves endoreplication that is followed by meiosis. We, therefore, tested whether Dmc1, a key component of the meiotic machinery, is required for pseudosexual reproduction. The meiotic recombinase gene *DMC1* was deleted in congenic strains H99α, VYD135α, and KN99**a,** and the resulting mutants were subjected to mating. A previous report documented that *dmc1*Δ bilateral crosses (both the parents are mutant for *DMC1*) display significantly reduced, but not completely abolished, sporulation in *Cryptococcus* (Lin et al., 2005). We observed a similar phenotype with the H99α *dmc1*Δ × KN99**a** *dmc1*Δ cross. While most of the basidia were devoid of spore chains, a small percent (21/760=2.7%) of the population bypassed the requirement for Dmc1 and produced spores (Figure 5A and S11A). When dissected, the germination frequency for these spores was found to be very low (~22% on average) with spores from many basidia not germinating at all (Table S3). Furthermore, *MAT*-specific PCRs revealed that some of the progeny were aneuploid or diploid. For VYD135α *dmc1*Δ × KN99**a** *dmc1*Δ, many fewer basidia (~0.1%) produced spore chains as compared to ~1% sporulation in VYD135α × KN99**a** (Figure 5A, B and S11B). *dmc1* mutant unilateral crosses (one of the two parents is mutant and the other one is wild-type) sporulated at a frequency of 0.4% suggesting that only one of the parental strains was producing spores (Figure 5B). When a few sporulating basidia from multiple mating spots of the VYD135α *dmc1*Δ × KN99**a** *dmc1*Δ bilateral cross were dissected, two different populations of basidia emerged, one with no spore germination, and the other with a high spore germination frequency and uniparental *MAT* inheritance (Table S3). We think that the basidia with a high spore germination frequency may represent examples that in some fashion have escaped the normal requirement for Dmc1. Combined together, the *DMC1* deletion led to a 20-fold reduction in viable sporulation in VYD135α × KN99**a** cross, observed as a 10-fold decrease from sporulation events in the bilateral cross and a further 2-fold reduction in the number of basidia producing viable spores.

**Figure 5.**
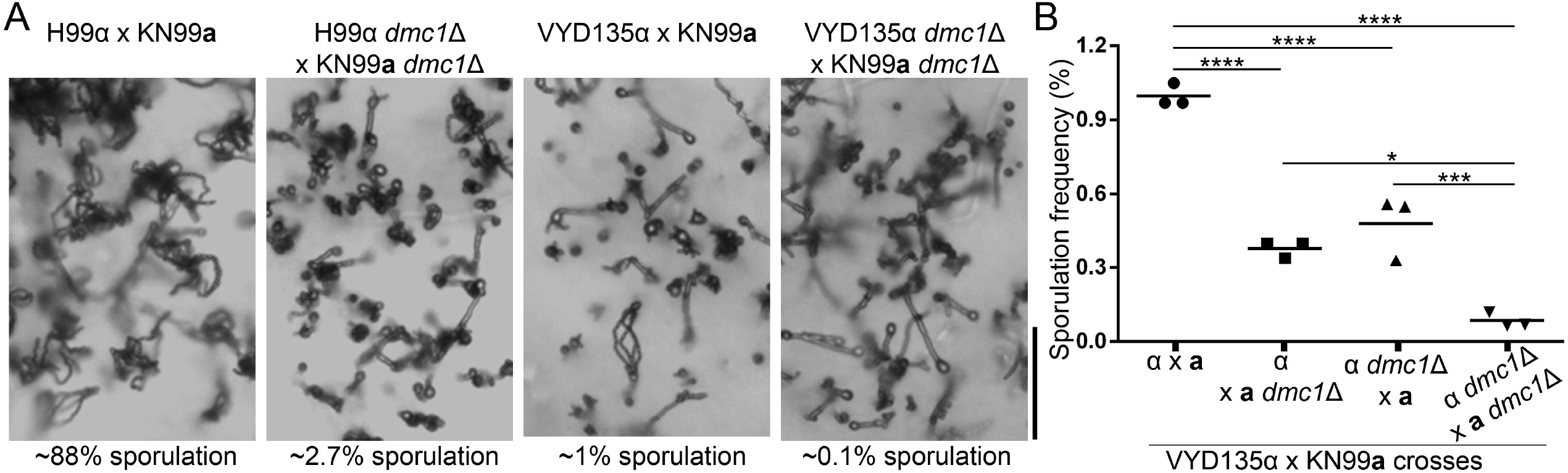
Meiotic recombinase Dmc1 is required for pseudosexual reproduction. **(A)** Light microscopy images showing the impact of *dmc1* mutation on sexual and pseudosexual reproduction in *C. neoformans*. Bar, 100 μm. **(B)** A graph showing quantification (n=3) of sporulation events in multiple crosses with *dmc1*Δ mutants. At least 3000 basidia were counted in each experiment.

To further support these conclusions, *DMC1* was deleted in mCherry-H4 tagged KN99**a** and crossed with GFP-H4 tagged VYD135α. We hypothesized that GFP-H4 tagged VYD135α would produce spore chains in this cross because it harbors *DMC1* whereas mCherry-H4 tagged KN99**a** *dmc1*Δ would fail to do so. Indeed, all 11 observed basidia with only the GFP-H4 fluorescence signal were found to produce spores but only 2 out of 19 mCherry-H4 containing basidia exhibited sporulation (Figure S12). These results combined with the spore dissection findings show that Dmc1 is critical for pseudosexual reproduction. While these results provide concrete evidence for meiosis as a part of pseudosexual reproduction, they also suggest the occurrence of a preceding endoreplication event. However, further studies will need to be conducted to validate and confirm endoreplication or alternate mechanisms.

## Discussion

Hybridogenesis and parthenogenesis are mechanisms that allow some organisms to overcome some hurdles of sexual reproduction and produce hemiclonal or clonal progeny (Avise, 2015; Hörandl, 2009; Lavanchy & Schwander, 2019). However, harmful mutations are not filtered in these processes, making them disadvantageous during evolution and thus restricting the occurrence of these processes to a limited number of animal species (Lavanchy & Schwander, 2019). In this study, we discovered and characterized the occurrence of a phenomenon in fungi that resembles hybridogenesis and termed it pseudosexual reproduction (Figure 6A). Fungi are known to exhibit asexual, (bi)sexual, unisexual, and parasexual reproduction and can switch between these reproductive modes depending on environmental conditions (Heitman, 2015; Heitman et al., 2013). The discovery of pseudosexual reproduction further diversifies known reproductive modes in fungi, suggesting the presence of sexual parasitism in this kingdom.

**Figure 6.**
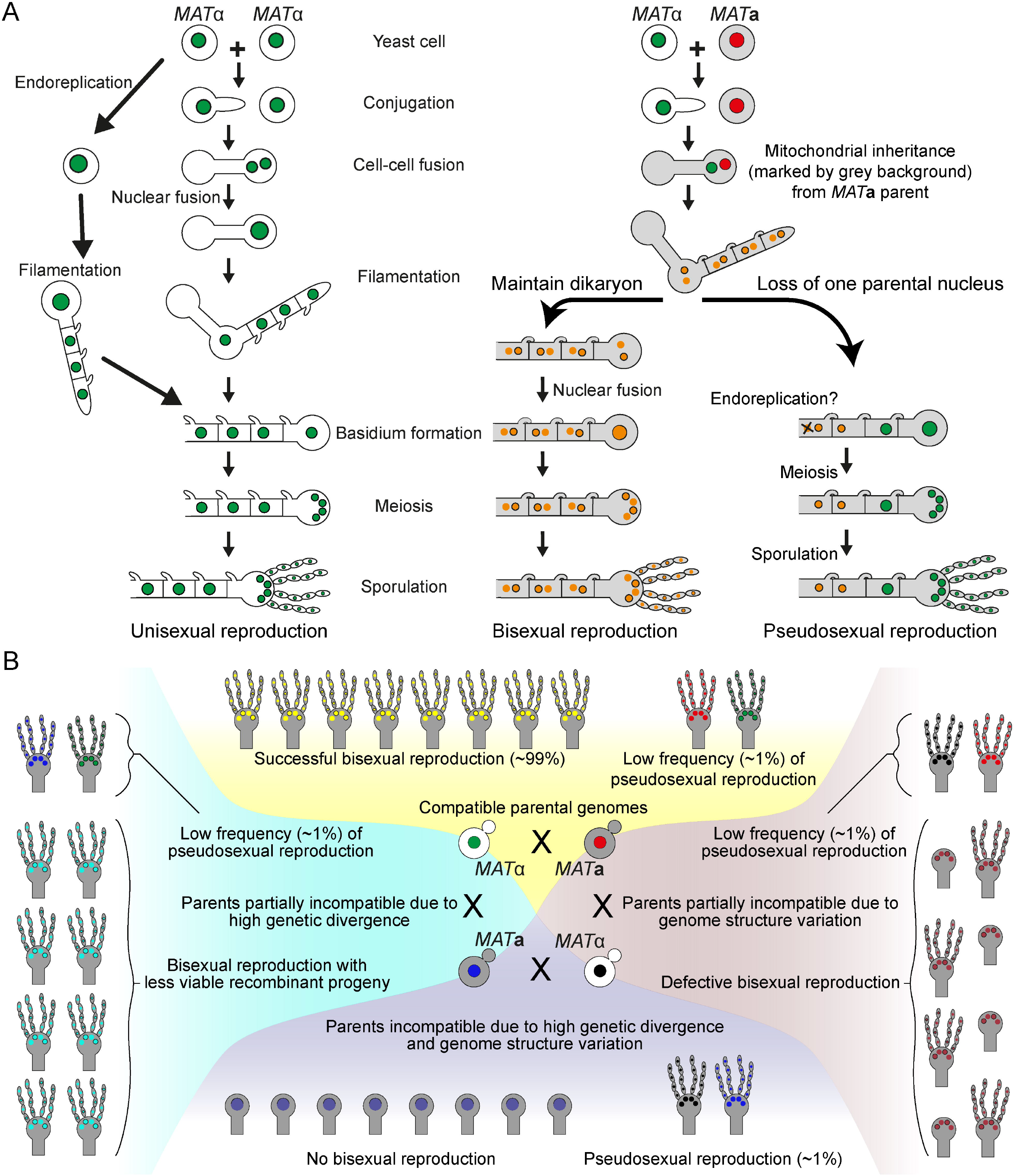
Occurrence of pseudosexual reproduction in *C. neoformans*. **(A)** A cartoon depicting various types of sexual reproduction in *Cryptococcus* species. *C. deneoformans* exhibits unisexual reproduction where two cells of the same mating-type fuse or a single cell undergo endoreplication followed by the production of clonal progeny. Both *C. neoformans* and *C. deneoformans* show bisexual reproduction where two cells of opposite mating-types fuse with each other and produce recombinant progeny. Pseudosexual reproduction, as observed in this study, arises from bisexual mating but generates clonal progeny for one of the parents after the other parental nucleus is lost during development. While both unisexual and pseudosexual reproduction produces clonal progeny, they differ with respect to the inheritance of mitochondrial DNA (marked by grey color cell background in the figure). **(B)** Scenarios showing the occurrence of pseudosexual reproduction under various hypothetical mating conditions. Except for one condition where the two parents are completely compatible with each other, pseudosexual reproduction could play a significant role in survival and dissemination despite its occurrence at a low frequency.

Hybridogenesis in animals occurs between two different species. The result of hybridogenesis is the production of gametes that are clones of one of the parents, which then fuse with an opposite-sex gamete of the second species, generating hemiclonal offspring. In our study, we observed a similar phenomenon where only one parent contributes to spores, the counterpart of mammalian gametes. However, we observed this phenomenon occurring between different strains of the same species, *C. neoformans*. It is important to note that these strains vary significantly from each other in terms of genetic divergence and in one case by chromosome rearrangements to the extent that they could be considered different species. This suggests that hybridogenesis in animals and pseudosexual reproduction in fungi are similar to each other. Hybridogenesis requires the exclusion of one of the parents, which is followed by endoreplication of the other parent’s genome and meiosis. The whole-genome sequence of the progeny in our study revealed the complete absence of one parent’s genome, suggesting manifestations of genome exclusion during hyphal growth. The mechanism by which the retained parental genome increases its ploidy before meiosis remains to be further investigated in *C. neoformans*. Endoreplication is known to occur in the sister species *C. deneoformans* during unisexual reproduction and we think that this is the most likely route via which ploidy is increased during pseudosexual reproduction.

The mechanism and time of genome exclusion during hybridogenesis in animals are not entirely understood, except for a few insights from triploid fishes of the genus *Poeciliopsis* and water frogs, *Pelophylax esculentus*. Studies using *Poeciliopsis* fishes showed that haploid paternal genome exclusion takes place during the onset of meiosis via the formation of a unipolar spindle, and thus only the diploid set of maternal chromosomes is retained (Cimino, 1972a, 1972b). On the other hand, studies involving *P. esculentus* revealed that genome exclusion occurs during mitotic division, before meiosis, which is followed by endoreplication of the other parental genome (Heppich, Tunner, & Greilhuber, 1982; Tunner & Heppich-Tunner, 1991; Tunner & Heppich, 1981). A recent study, however, proposed that genome exclusion in *P. esculentus* could also take place during early meiotic phases (Dolezalkova et al., 2016). Using fluorescence microscopy, we examined the steps of nuclear exclusion in *C. neoformans* and found that it occurs during mitotic hyphal growth and not during meiosis. We also observed that genome exclusion could happen with either of the two parents in *C. neoformans*, similar to what has also been reported for water frogs. However, for most other species, genome exclusion was found to occur with the male genome only, leaving behind the female genome for meiosis (Cimino, 1972a; Holsbeek & Jooris, 2009; Lavanchy & Schwander, 2019; Umphrey, 2006; Uzzell, Günther, & Berger, 1976; Vinogradov, Borkin, Gunther, & Rosanov, 1991). Multiple studies have showed the formation of meiotic synaptonemal complex during hybridogenesis clearly establishing the presence of meiosis during this process (Dedukh et al., 2019; Dedukh et al., 2020; Nabais, Pereira, Cunado, & Collares-Pereira, 2012). Our results showed that the meiotic recombinase Dmc1 is required for pseudosexual reproduction suggesting the presence of meiosis, whereas there is no direct evidence for the role of a meiotic recombinase in hybridogenetic animals. Taken together, these results indicate that the mechanism might be at least partially conserved across distantly related species. Future studies will shed more light on this and if established, the amenability of *C. neoformans* to genetic manipulation will aid in deciphering some of the unanswered questions related to hybridogenesis in animals.

The occurrence of pseudosexual reproduction might also have significant implications for *C. neoformans* biology. Most (>95%) of *Cryptococcus* natural isolates belong to only one mating type, α (Zhao et al., 2019). While the reason behind this distribution is unknown, one explanation could be the presence of unisexual reproduction in the sister species *C. deneoformans* and *C. gattii* (Fraser et al., 2005; Lin et al., 2005; Phadke, Feretzaki, Clancey, Mueller, & Heitman, 2014). The presence of pseudosexual reproduction in *C. neoformans* might help explain the mating-type distribution pattern for this species. In this report, one of the *MAT***a** natural isolates, Bt63**a**, did not contribute to pseudosexual reproduction and the other isolate, IUM96**a**, produced uniparental progeny in only one basidium, while the rest of the basidia produced *MAT*α progeny. We hypothesize that *MAT***a** isolates may be defective in this process due to either a variation in their genomes or some other as yet undefined sporulation factor. As a result, pseudosexual reproduction would result in the generation of predominantly α progeny in nature reducing the *MAT***a** population and thus favoring the expansion of the α mating-type population. Whether pseudosexual reproduction occurs in other pathogenic species such as *C. deneoformans* and non-pathogenic species such as *C. amylolentus* will be investigated in future studies. Attempts to identify the occurrence of pseudosexual reproduction between species where hybrids are known to occur, *C. neoformans* and *C. deneoformans* hybrids, will also be made. These studies will help establish the scope of pseudosexual reproduction in *Cryptococcus* species and could be extended to other basidiomycetes.

We propose that pseudosexual reproduction can occur between any two opposite mating-type strains as long as each of them is capable of undergoing cell-cell fusion and at least one of them can sporulate. We speculate that pseudosexual reproduction might play a key role in *C. neoformans* survival during unfavorable conditions. In conditions where two mating partners are fully compatible, pseudosexual reproduction will be mostly hidden and might not be important (Figure 6B, top panel). However, when the two mating partners are partially incompatible or completely incompatible due to high genetic divergence or karyotypic variation, pseudosexual reproduction will be important (Figure 6B, left, right, and bottom panels). For example, most of the basidia in H99α and Bt63**a** cross largely produce aneuploid and/or inviable progeny leading to unsuccessful sexual reproduction. However, a small yet significant proportion of the basidia generate clonal yet viable and fit progeny via pseudosexual reproduction. We hypothesize that these progeny will have a better chance of survival and find a suitable mating partner in the environment whereas the unfit recombinant progeny might fail to do so. In nature, this might allow a new genotype/karyotype to not only survive but also expand and will prove advantageous. If a new genotype/karyotype had only the option of undergoing sexual reproduction, it might not survive, restricting the evolution of a new strain. Overall, this mode of pseudosexual reproduction might act as an escape path from genomic incompatibilities between two related isolates and allow them to produce spores for dispersal and infection.

The fungal kingdom is one of the more diverse kingdoms with approximately 3 million species (Sun, Hoy, & Heitman, 2020). The finding of hybridogenesis-like pseudosexual reproduction hints towards unexplored biology in this kingdom that might provide crucial clues for understanding the evolution of sex. Fungi have also been the basis of studies focused on understanding the evolution of meiosis, and the presence of genome reduction, as well as the parasexual cycle in fungi, have led to the proposal that meiosis evolved from mitosis (Hurst & Nurse, 1991; Wilkins & Holliday, 2009). Pseudosexual reproduction may be a part of an evolutionary process wherein genome exclusion followed by endoreplication and meiosis was an ancestral form of reproduction that preceded the evolution of sexual reproduction. Evidence supporting such a hypothesis can be observed in organisms undergoing facultative sex or facultative parthenogenesis (Booth et al., 2012; Fields, Feldheim, Poulakis, & Chapman, 2015; Hodač, Klatt, Hojsgaard, Sharbel, & Hörandl, 2019; Hojsgaard & Harandl, 2015). The presence of these organisms also suggests that a combination of both sexual and clonal modes of reproduction might prove to be evolutionarily advantageous.

## Materials and Methods

### Strains and media

*C. neoformans* wild-type strains H99α and KN99**a** served as the wild-type isogenic parental lineages for the experiments, in addition to *MAT***a** strains Bt63**a** and IUM96-2828**a**. Strains were grown in YPD media for all experiments at 30°C unless stated otherwise. G418 and/or NAT were added at a final concentration of 200 and 100 μg/ml, respectively, for the selection of transformants. MS media was used for all the mating assays, which were performed as described previously (Sun, Priest, et al., 2019). Basidia-specific spore dissections were performed after two-five weeks of mating, and the spore germination frequency was scored after five days of dissection. All strains and primers used in this study are listed in Table S4 and S5, respectively.

### Genotyping for mating-type locus and mitochondria

Mating-type (*MAT*) and mitochondrial genotyping for all the progeny were conducted using PCR assays. Genomic DNA was prepared using the MasterPure™ Yeast DNA purification kit from Lucigen. To determine the *MAT*, the *STE20* allele present within the *MAT* locus was detected since it differs in length between the two different mating type strains. Primers specific to both *MAT***a** and *MAT*α (JOHE50979-50982 in Table S5) were mixed in the same PCR mix and the identification was made based on the length of the amplicon (Figure 1E–G). For the mitochondrial genotyping, the *COX1* allele present in the mitochondrial DNA was probed to distinguish between H99α/VYD135α and KN99**a**/IUM96**a**. For the differentiation between Bt63**a** and H99α/VYD135α, the *COB1* allele was used because *COX1* in Bt63**a** is identical to H99α/VYD135α. The difference for both *COX1* and *COB1* is the presence or absence of an intron and results in significantly different size products between *MAT*α and *MAT***a** parents (Figure 1 and S3). The primers used for these assays (JOHE51004-51007) are mentioned in Table S5.

### Genomic DNA isolation for sequencing

Genomic DNA for whole-genome sequencing was prepared using the CTAB-based lysis method, as described previously (Yadav et al., 2020). Briefly, 50 ml of an overnight culture was pelleted, frozen at −80°C, and subjected to lyophilization. The lyophilized cell pellet was broken into a fine powder, mixed with lysis buffer, and the mix was incubated at 65°C for an hour with intermittent shaking. The mix was then cooled on ice, and the supernatant was transferred into a fresh tube, and an equal volume of chloroform (~15 ml) was added and mixed. The mix was centrifuged at 3200 rpm for 10 min, and the supernatant was transferred to a fresh tube. An equal volume of isopropanol (~18 to 20 ml) was added into the supernatant and mixed gently. This mix was incubated at −20°C for an hour and centrifuged at 3200 rpm for 10 min. The supernatant was discarded, and the DNA pellet was washed with 70% ethanol. The pellet was air-dried and dissolved in 1ml of RNase containing 1X TE buffer and incubated at 37°C for 45 min. The DNA was again chloroform purified and precipitated using isopropanol, followed by ethanol washing, air drying, and finally dissolved in 200 μl 1X TE buffer. The DNA quality was estimated with NanoDrop, whereas DNA quantity was estimated with Qubit.

### Whole-genome Illumina sequencing, ploidy, and SNP analysis

Illumina sequencing of the strains was performed at the Duke sequencing facility core (https://genome.duke.edu/), using Novaseq 6000 as 150 paired-end sequencing. The Illumina reads, thus obtained, were mapped to the respective genome assembly (H99, VYD135, Bt63, or IUM96) using Geneious default mapper to estimate ploidy. The resulting BAM file was converted to a .tdf file, which was then visualized through IGV to estimate the ploidy based on read coverage for each chromosome.

For SNP calling and score for recombination in the progeny, Illumina sequencing data for each progeny was mapped to parental strain genome assemblies individually using the Geneious default mapper with three iterations. The mapped BAM files were used to perform variant calling using Geneious with 0.8 variant frequency parameter and at least 90x coverage for each variant. The variants thus called were exported as VCF files and imported into IGV for visualization purposes. H99, Bt63, IUM96-2828, and VYD135 Illumina reads were used as controls for SNP calling analysis.

### PacBio/Nanopore genome assembly and synteny comparison

To obtain high-molecular-weight DNA for Bt63 genome PacBio and IUM96-2828 genome Nanopore sequencing, DNA was prepared as described above. The size estimation of DNA was carried out by electrophoresis of DNA samples using PFGE. For this purpose, the PFGE was carried out at 6V/cm at a switching frequency of 1 to 6 sec for 16 h at 14°C. Samples with most of the DNA ≥100 kb or larger were selected for sequencing. For PacBio sequencing, the DNA sample was submitted to the Duke sequencing facility core. Nanopore sequencing was performed in our lab using a MinION device on an R9.4.1 flow cell. After sequencing, reads were assembled to obtain a Bt63 genome assembly via Canu using PacBio reads > 2 kb followed by five rounds of pilon polishing. For IUM96-2828, one round of nanopolish was also performed before pilon polishing. Once completed, the chromosomes were numbered based on their synteny with the H99 genome. For chromosomes involved in translocation (Chr 3 and Chr 11), the chromosome numbering was defined by the presence of the respective syntenic centromere from H99. Centromere locations were mapped based on BLASTn analysis with H99 centromere flanking genes.

Synteny comparisons between the genomes were performed with SyMAP v4.2 using default parameters (Soderlund, Bomhoff, & Nelson, 2011) (http://www.agcol.arizona.edu/software/symap/). The comparison block maps were exported as .svg files and were then processed using Adobe^®^ Illustrator^®^ and Adobe^®^ Photoshop^®^ for representation purposes. The H99 genome was used as the reference for comparison purposes for plotting VYD135, Bt63, and IUM96-2828 genomes. The centromere and telomere locations were manually added during the figure processing.

### Fluorescent tagging and microscopy

GFP and mCherry tagging of histone H4 were performed by integrating respective constructs at the safe haven locus (Arras, Chitty, Blake, Schulz, & Fraser, 2015). GFP-H4 tagging was done using the previously described construct, pVY3 (Yadav & Sanyal, 2018). For mCherry-H4 tagging, the GFP-containing fragment in pVY3 was excised using SacI and BamHI and was replaced with mCherry sequence PCR amplified from the plasmid pLKB25 (Kozubowski & Heitman, 2010). The constructs were then linearized using XmnI and transformed into desired strains using CRISPR transformation, as described previously (Fan & Lin, 2018). The transformants were screened by PCR, and correct integrants were obtained and verified using fluorescent microscopy.

To observe the fluorescence signals in the hyphae and basidia, a 2-3 week old mating patch was cut out of the plate and directly inverted onto a coverslip in a glass-bottom dish. The dish was then used to observe filaments under a DeltaVision microscope available at the Duke University Light Microscopy Core Facility (https://microscopy.duke.edu/dv). The images were captured at 60X magnification with 2×2 bin size and z-sections of either 1 or 0.4 μm each. GFP and mCherry signals were captured using the GFP and mCherry filters in the Live-Cell filter set. The images were processed using Fiji-ImageJ (https://imagej.net/Fiji) and exported as tiff files as individual maximum projected images. The final figure was then assembled using Adobe^®^ Photoshop^®^ software for quality purposes.

### Sporulation frequency counting

To visualize hyphal growth and sporulation defects during mating assays, the mating plates were directly observed under a Nikon Eclipse E400 microscope. Hyphal growth and basidia images were captured using the top-mounted Nikon DXM1200F camera on the microscope. The images were processed using Fiji-ImageJ and assembled in Adobe^®^ Photoshop^®^ software.

For crosses involving wild-type H99α, VYD135α, KN99**a**, Bt63**a**, IUM96**a**, approximately 1000 total basidia were counted after 4 weeks of mating, and the sporulation frequency was calculated. For crosses involving VYD135 *dmc1*Δ strain, three mating spots were setup independently. From each mating spot periphery, 6 images were captured after 3-4 weeks of mating. Basidia (both sporulating and non-sporulating) in each of these spots were counted manually after some processing of images using ImageJ. The sporulation frequency was determined by dividing the sporulating basidia by the total number of basidia for each spot. Each mating spot was considered as an independent experiment and at least 3000 basidia were counted from each mating spot.

### Flow cytometry

Flow cytometry analysis was performed as described previously (Fu & Heitman, 2017). Cells were grown on YPD medium for two days at 30°C, harvested, and washed with 1X PBS buffer followed by fixation in 70% ethanol at 4°C overnight. Next, cells were washed once with 1 ml of NS buffer (10 mM Tris-HCl, pH = 7.2, 250 mM sucrose, 1 mM EDTA, pH = 8.0, 1 mM MgCl2, 0.1 mM CaCl2, 0.1 mM ZnCl2, 0.4 mM phenylmethylsulfonyl fluoride, and 7 mM β-mercaptoethanol), and finally resuspended in 180 μl NS buffer containing 20 μl 10 mg/ml RNase and 5 μl 0.5 mg/ml propidium iodide (PI) at 37°C for 3-4 hours. Then, 50 μl stained cells were diluted in 2 ml of 50 mM Tris-HCl, pH = 8.0, transferred to FACS compatible tube, and submitted for analysis at the Duke Cancer Institute Flow Cytometry Shared Resource. For each sample, 10,000 cells were analyzed on the FL1 channel on the Becton-Dickinson FACScan. Wild-type H99 and previously generated AI187 were used as haploid and diploid controls, respectively, in these experiments. Data analysis was performed using the FlowJo software.

### Data Availability

The sequence data generated in this study were submitted to NCBI with the BioProject accession number PRJNA682203.

## Acknowledgments

We thank Shelby Priest and Arti Dumbrepatil for critical reading of this manuscript. This work was supported by NIH/NIAID R37 MERIT award AI39115-24, R01 grant AI50113-16 awarded to JH; and R01 grant AI33654-04 awarded to JH, David Tobin, and Paul Magwene. JH is also Co-Director and Fellow of the CIFAR program *Fungal Kingdom: Threats & Opportunities*.

## Competing interests

The authors declare no competing interests.

## Notes

### Competing Interest Statement

The authors have declared no competing interest.

### Summary of Updates

We termed the process described as pseudosexual reproduction reflecting the finding that although the observed process starts with bisexual mating, the two parental nuclei do not recombine and the progeny are clonal with the nuclear genome that is identical to one of the two parents. We added additional data definitively demonstrating mitochondrial uniparental inheritance during pseudosexual reproduction by both PCR as well as whole-genome sequencing.

