## Supplementary material for "Uniparental nuclear inheritance following bisexual mating in fungi": Suplementary Information

for

### Supplementary Tables

#### Table S1. The genotype of basidia-specific spores dissected from H99α x Bt63a and VYD135α x Bt63a crosses.

| | Basidia # | Spores dissected | Spores germinated | % germinated | *MAT* | Mitochondria |
|---|---|---|---|---|---|---|
| **H99α x Bt63a cross (2 weeks old)** | 1 | 14 | 2 | 14 | 1**a** + 1**a**/α | **a** |
| | 2 | 14 | 5 | 36 | 3α + 1**a** + 1**a**/α | **a** |
| | 3 | 14 | 2 | 14 | 1α + 1**a** | **a** |
| | 4 | 14 | 0 | 0 | NA | NA |
| | 5 | 14 | 3 | 21 | 1a + 2**a**/α | **a** |
| | 6 | 14 | 13 | 93 | 4α + 5**a** + 4**a**/α | **a** |
| | 7 | 14 | 3 | 21 | 2α + 1**a**/α | **a** |
| | 8 | 9 | 0 | 0 | NA | NA |
| | 9 | 12 | 3 | 25 | 1α + 2**a** | **a** |
| | 10 | 22 | 6 | 27 | 5α + 1**a**/α | **a** |
| **VYD135α x Bt63a cross (2 weeks old)** | 1 | 6 | 0 | 0 | NA | NA |
| | 2 | 14 | 6 | 43 | All α | **a** |
| | 3 | 13 | 13 | 100 | All α | **a** |
| | 4 | 26 | 0 | 0 | NA | NA |
| | 5 | 10 | 7 | 70 | All α | **a** |
| | 6 | 14 | 0 | 0 | NA | NA |
| | 7 | 14 | 0 | 0 | NA | NA |
| | 8 | 14 | 13 | 93 | All α | **a** |
| | 9 | 20 | 18 | 90 | All α | **a** |
| | 10 | 16 | 0 | 0 | NA | NA |
| **VYD135α x Bt63a cross (5 weeks old)** | 11 | 14 | 11 | 79 | All α | **a** |
| | 12 | 14 | 6 | 43 | All α | **a** |
| | 13 | 28 | 18 | 64 | All α | **a** |
| | 14 | 14 | 12 | 86 | All α | **a** |
| | 15 | 7 | 4 | 57 | All α | **a** |
| | 16 | 12 | 9 | 75 | All α | **a** |
| | 17 | 28 | 19 | 68 | All α | **a** |
| | 18 | 22 | 6 | 27 | All α | **a** |
| | 19 | 14 | 7 | 50 | All α | **a** |
| | 20 | 12 | 12 | 100 | All α | **a** |

NA – Not Applicable

**Table S2. The genotype of basidia-specific spores dissected from H99α x IUM96-2828a and VYD135α x IUM96-2828a crosses.**

| | Basidia # | Spores dissected | Spores germinated | % germinated | *MAT* | Mitochondria |
|---|---|---|---|---|---|---|
| **H99α x IUM96-2828a cross (2 weeks old)** | 1 | 26 | 21 | 81 | 10α + 11**a** | **a** |
| | 2 | 28 | 28 | 100 | 16α + 12**a** | **a** |
| | 3 | 14 | 9 | 64 | 6α + 3**a** | **a** |
| | 4 | 20 | 6 | 30 | 4α + 1**a** + 1**a**/α | **a** |
| | 5 | 22 | 0 | 0 | NA | NA |
| | 6 | 15 | 0 | 0 | NA | NA |
| **H99α x IUM96-2828a cross (5 weeks old)** | 7 | 19 | 17 | 89 | All α | **a** |
| | 8 | 20 | 20 | 100 | All α | **a** |
| | 9 | 20 | 18 | 90 | All α | **a** |
| | 10 | 26 | 7 | 27 | **a**/α | **a** |
| | 11 | 14 | 2 | 14 | All α | **a** |
| | 12 | 14 | 5 | 36 | 3α + 1**a** + 1**a**/α | **a** |
| **VYD135α x IUM96-2828a cross (2 weeks old)** | 1 | 12 | 0 | 0 | NA | NA |
| | 2 | 23 | 0 | 0 | NA | NA |
| | 3 | 14 | 11 | 79 | All α | **a** |
| | 4 | 13 | 0 | 0 | NA | NA |
| | 5 | 16 | 3 | 19 | All **a** | **a** |
| | 6 | 14 | 0 | 0 | NA | NA |
| **VYD135α x IUM96-2828a cross (5 weeks old)** | 7 | 14 | 0 | 0 | All α | **a** |
| | 8 | 23 | 10 | 43 | All α | **a** |
| | 9 | 22 | 19 | 86 | All α | **a** |
| | 10 | 23 | 22 | 96 | All α | **a** |
| | 11 | 28 | 24 | 86 | All α | **a** |
| | 12 | 28 | 14 | 50 | All α | **a** |
| | 13 | 22 | 20 | 91 | All α | α |
| | 14 | 22 | 22 | 100 | All α | **a** |
| | 15 | 24 | 13 | 54 | All α | **a** |
| | 16 | 17 | 8 | 47 | All α | **a** |
| | 17 | 9 | 8 | 89 | All α | **a** |
| | 18 | 24 | 9 | 38 | All α | **a** |

NA – Not Applicable

**Table S3. Genotype analysis of basidia-specific progeny from H99α *dmc1*Δ x KN99a *dmc1*Δ and VYD135α *dmc1*Δ x KN99a *dmc1*Δ crosses.**

| **Basidia #** | H99α *dmc1*Δ x KN99**a** *dmc1*Δ cross | | | | VYD135α *dmc1*Δ x KN99a *dmc1*Δ cross | | | |
|---|---|---|---|---|---|---|---|---|
| | **Spores germinated/ dissected** | **% germinated** | ***MAT*** | **Mito** | **Spores germinated/ dissected** | **% germinated** | ***MAT*** | **Mito** |
| 1 | 7/24 | 29 | 2α + 5**a**/α | **a** | 0/26 | 0 | **-** | **-** |
| 2 | 2/20 | 10 | All α | **a** | 7/14 | 50 | All **a** | **a** |
| 3 | 3/20 | 15 | All α | **a** | 0/10 | 0 | **-** | **-** |
| 4 | 5/14 | 36 | All **a**/α | **a** | 8/18 | 44 | All **a** | **a** |
| 5 | 3/11 | 27 | All **a** | **a** | 12/12 | 100 | All **a** | **a** |
| 6 | 0/12 | 0 | - | **-** | 7/8 | 88 | All **a** | **a** |
| 7 | 7/26 | 27 | All **a** | **a** | 14/14 | 100 | All **a** | **a** |
| 8 | 0/9 | 0 | - | **-** | 0/8 | 0 | **-** | **-** |
| 9 | 0/19 | 0 | - | **-** | 0/14 | 0 | **-** | **-** |
| 10 | 1/11 | 9 | α | **a** | 19/19 | 100 | All **a** | **a** |
| 11 | 24/27 | 89 | 12α + 6**a** + 6**a**/α | **a** | 0/5 | 0 | **-** | **-** |
| 12 | 5/22 | 23 | 1α + 4**a**/α | **a** | - | - | **-** | **-** |

Mito refers to Mitochondria.

**Table S4. Strains used in this study.**

| Strain name | Description | Reference |
|---|---|---|
| H99 | Wild-type *MAT*α | (Perfect, Ketabchi, Cox, Ingram, & Beiser, 1993) |
| KN99**a** | Wild-type *MAT***a** | (Nielsen et al., 2003) |
| VYD135 | H99 derivative with shuffled chromosomes | (Yadav, Sun, Coelho, & Heitman, 2020) |
| Bt63 | Wild-type *MAT***a** | (Litvintseva et al., 2003) |
| IUM96-2828 | Wild-type *MAT***a** | (Keller, Viviani, Esposto, Cogliati, & Wickes, 2003) |
| VYD158 | H99α *GFP-H4*::*NAT* | This study |
| VYD159 | VYD135α *GFP-H4*::*NAT* | This study |
| VYD160 | KN99**a** *mCherry-H4*::*NAT* | This study |
| VYD171 | Bt63**a** *mCherry-H4*::*NAT* | This study |
| VYD178 | H99α *dmc1*Δ::*NEO* | This study |
| VYD179 | KN99a *dmc1*Δ::*NEO* | This study |
| VYD181 | VYD135 *dmc1*Δ::*NEO* | This study |
| VYD182 | KN99**a** *mCherry-H4*::*NAT* *dmc1*Δ::*NEO* | This study |

**Table S5. Primers used in this study.**

| Primer # | Sequence | Purpose |
|---|---|---|
| JOHE50979 | CTAACTCTACTACACCTCACGGCA | *MAT***a** PCR |
| JOHE50980 | CGCACTGCAAAATAGATAAGTCTG | |
| JOHE50981 | GGCTGCAATCACAGCACCTTAC | *MAT*α PCR |
| JOHE50982 | CTTCATGACATCACTCCCCTAT | |
| JOHE51004 | TGGTGGTGGTGACCCAGTTCT | Mitochondria genotyping primers for H99, VYD135, KN99, IUM96 |
| JOHE51005 | CCGAAGATCTTAGGTGCCCA | |
| JOHE51006 | CCACAACCTATTAACATTAGCTACGC | Mitochondria genotyping primers for H99, VYD135, Bt63 |
| JOHE51007 | CGTCTCCATCTACAAAGCCAGCAAAC | |
| JOHE51159 | ACCGGCAGGGTATACTGTTGAGTGCTGTGGTGAAAG AGATGTTTTAGAGCTAGAAATAGC | Safe Haven guide RNA |
| JOHE51204 | GCATGCGAGCTCGGCAGATACGATATGTTGGCG | $P_{H3}$-mCherry contruct |
| JOHE51205 | CCTCCTCGCCCTTGCTCACCATTGATAGATGTGTTGT GGTG | |
| JOHE51206 | CACCACAACACATCTATCAATGGTGAGCAAGGGCGA GGAGG | |
| JOHE51207 | GCATGCGGATCCCTTGTACAGCTCGTCCATGCC | |
| JOHE51403 | GTCCAAAGTACTCAATGATACGACC | Dmc1 deletion construct |
| JOHE51404 | CAGCTCACATCCTCGCAGCCATAGGAAGGGTGGCAA ATCCAAATGAC | |
| JOHE51405 | GTCATTTGGATTTGCCACCCTTCCTATGGCTGCGAGG ATGTGAGCTG | |
| JOHE51406 | GTCGAGTCAAAATGAATAGGAATCCGCGGTTTATCT GTATTAACACGG | |
| JOHE51407 | CCGTGTTAATACAGATAAACCGCGGATTCCTATTCAT TTTGACTCGAC | |
| JOHE51408 | AATTGTGGAACAAGAACGAGGAC | |
| JOHE51409 | TAGCCAACATACCCAAACCACCAGC | |
| JOHE51410 | GTGGTTGTTTGAGTCGTTTGAGAGC | |
| JOHE51416 | ACCGGCAGGGTATACTGTTGATAATAGGCCGGCTTCT GGTGTTTTAGAGCTAGAAATAGC | Dmc1 deletion guide RNA |
| JOHE51417 | ACCGGCAGGGTATACTGTTGGATTCACCTGATATGCC GGAGTTTTAGAGCTAGAAATAGC | |

#### Supplementary Figures and Figure Legends

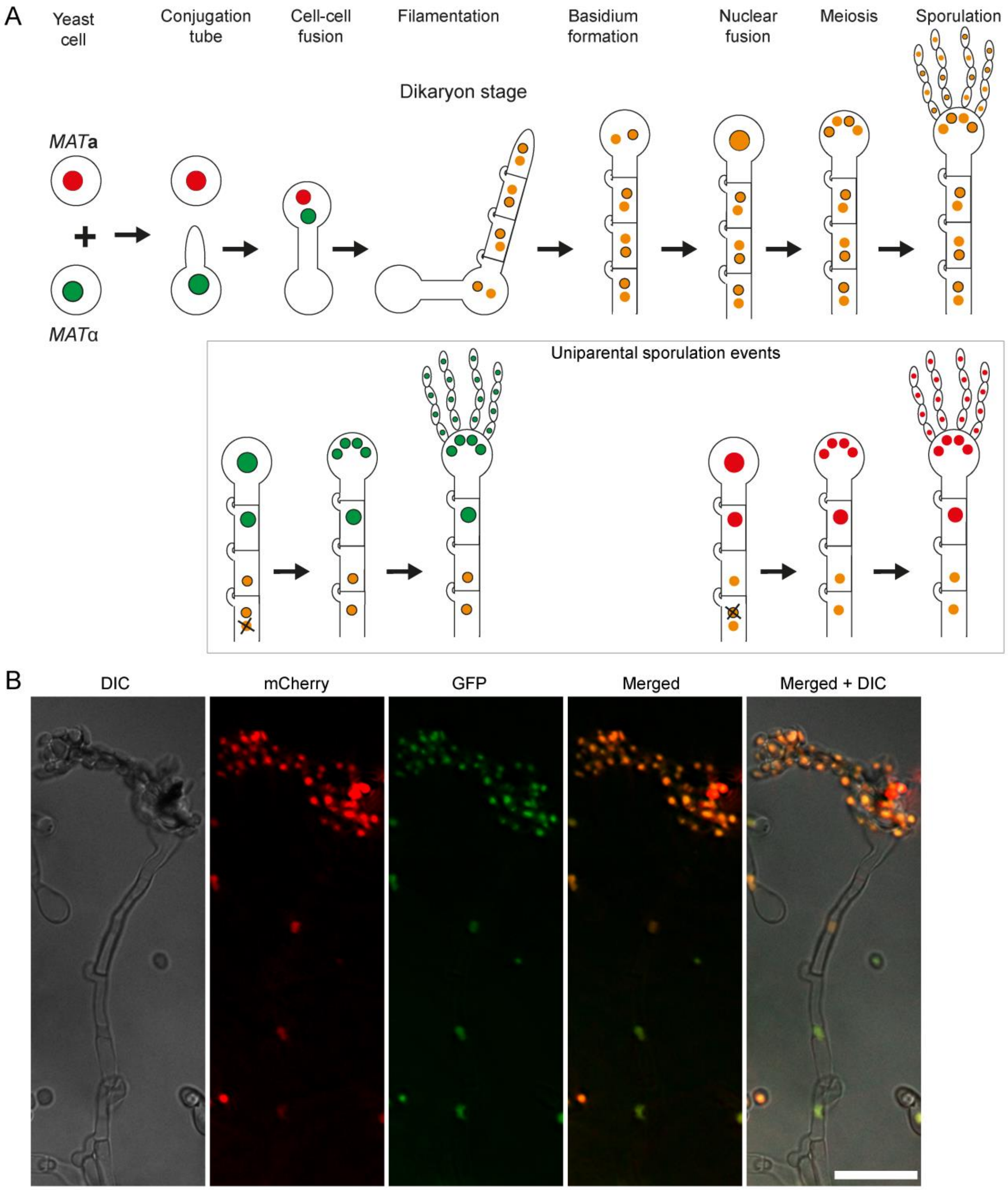


**Figure S1. Dynamics of sexual reproduction and sporulation analyzed with *C. neoformans* strains expressing nuclear-localized fluorescent reporter proteins. (A)** A cartoon depicting various stages of sexual reproduction in *C. neoformans*, from the formation of conjugation tube to sporulation, and possible dynamics of the nuclei at these different stages. After cell-cell fusion, tagged proteins assort into both nuclei and yield a yellow/orange fluorescence color as a result of the mixing of the green and red signals. Cartoons in the box show hypothetical scenarios where uniparental nuclear inheritance occurs after the loss of one parental nucleus. **(B)** Direct fluorescence microscopy images showing the status of GFP-H4 and mCherry-H4 tagged nuclei in post-mating hyphae as well as in spores. Both GFP and mCherry fluorescent colors were observed in hyphae and spores as hypothesized in A. Bar, 10 µm.

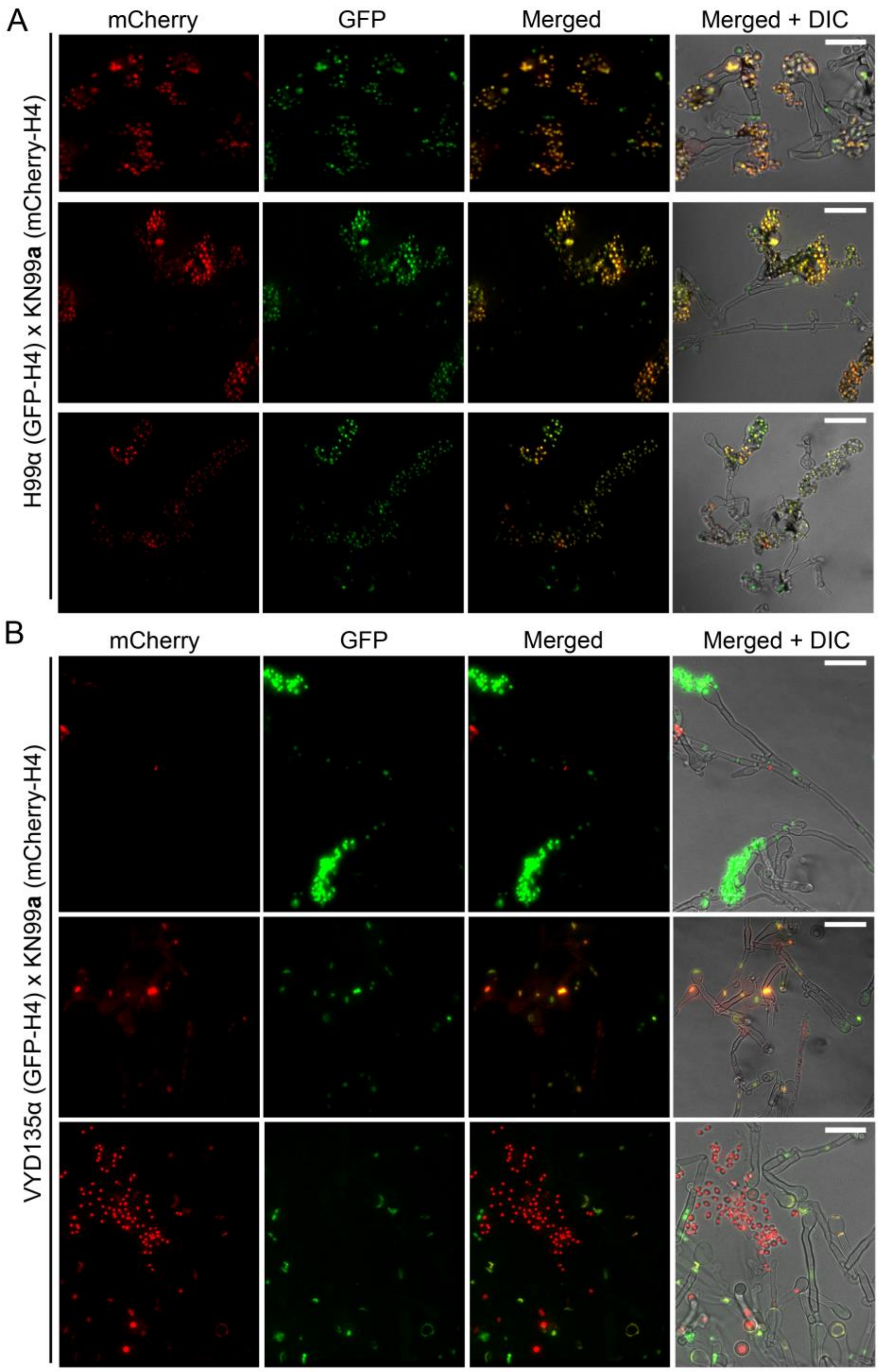


**Figure S2. Nuclear dynamics during sporulation in the wild-type and VYD135α crosses.** GFP-H4 and mCherry-H4 tagging revealed different localization patterns in the **(A)** wild-type H99α x KN99**a** and **(B)** VYD135α x KN99**a** crosses. Wild-type spore chains mostly harbored both the nuclear stains as a result of bisexual meiosis. On the other hand, basidia with only one of the parental nuclei produced spores in VYD135α x KN99**a** crosses; basidia with both nuclei failed to produce spore chains and, as a result, remained as bald basidia. Bars, 10 µm.

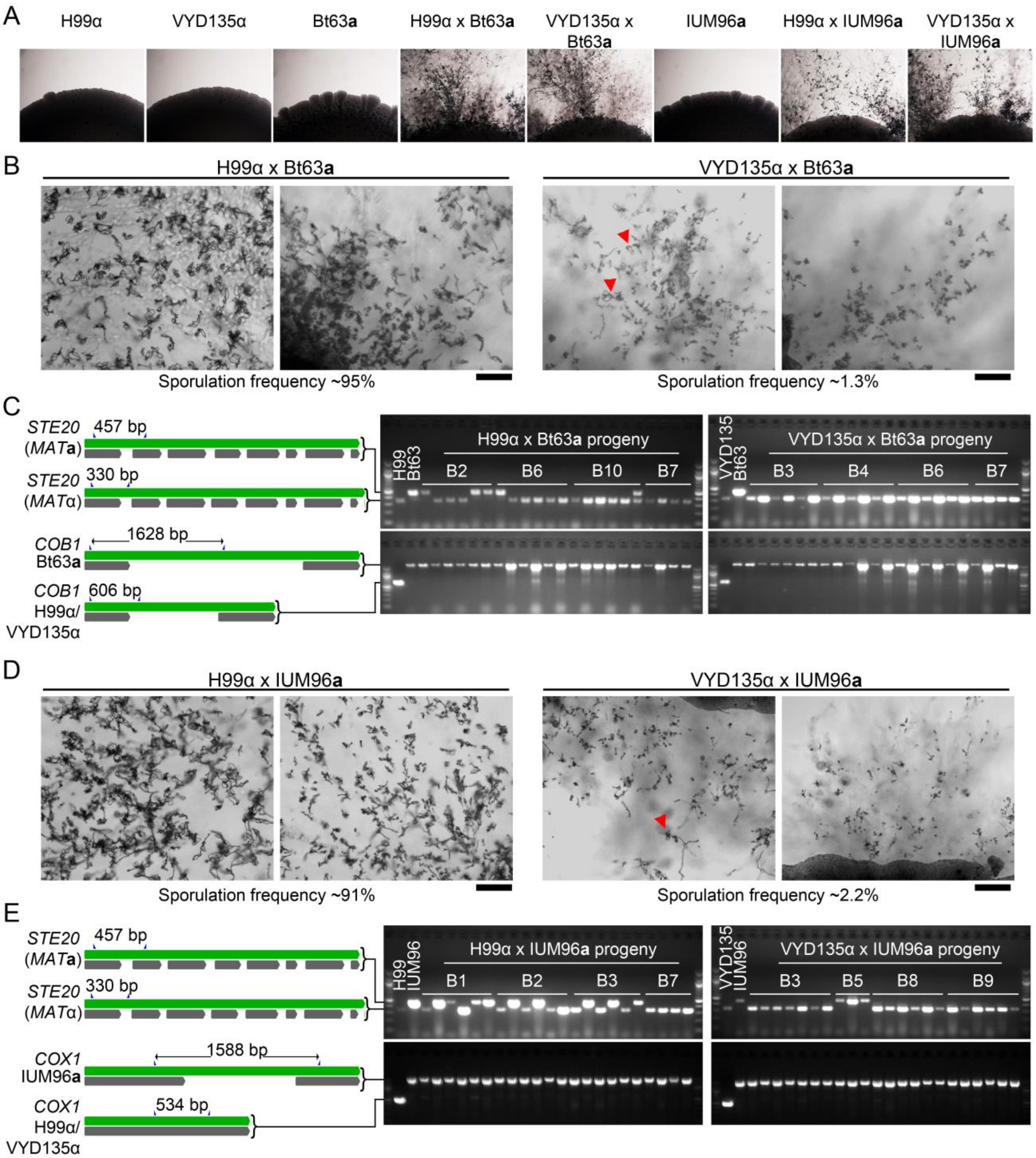


**Figure S3. Pseudosexual reproduction occurs in natural isolates, Bt63a and IUM96a. (A)** Images of the mating spots showing filamentation when two strains of opposite mating-type are crossed. No filamentation is observed without the presence of a mating partner. **(B and D)** Light microscopy images showing sporulation frequency in crosses involving Bt63**a** **(B)** and IUM96**a** **(D)**. Bars, 100 µm. **(C and E)** Schemes depicting the *STE20* alleles used for *MAT* locus and *COB1* (for Bt63**a**) and *COX1* (for IUM96**a**) alleles for mitochondrial genotyping, respectively.

Gel images show the PCR analysis on progeny from four basidia and the parental strains for all crosses as mentioned.

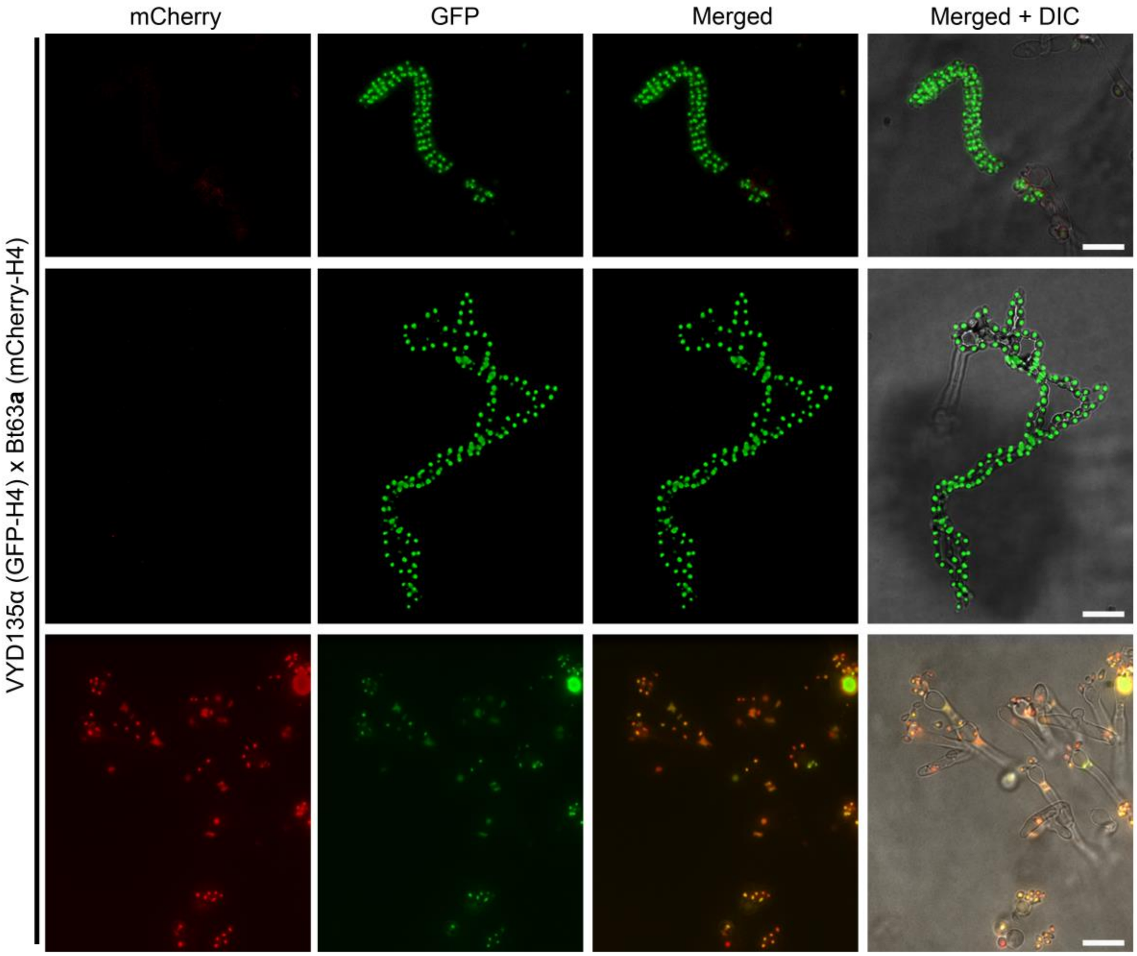


**Figure S4. Bt63a fluorescence microscopy revealed pseudosexual reproduction events.** GFP-H4 tagged VYD135α crossed with mCherry-H4 tagged Bt63**a** showed only VYD135α sporulation events as also observed in spore dissection analysis. Bars, 10 µm.

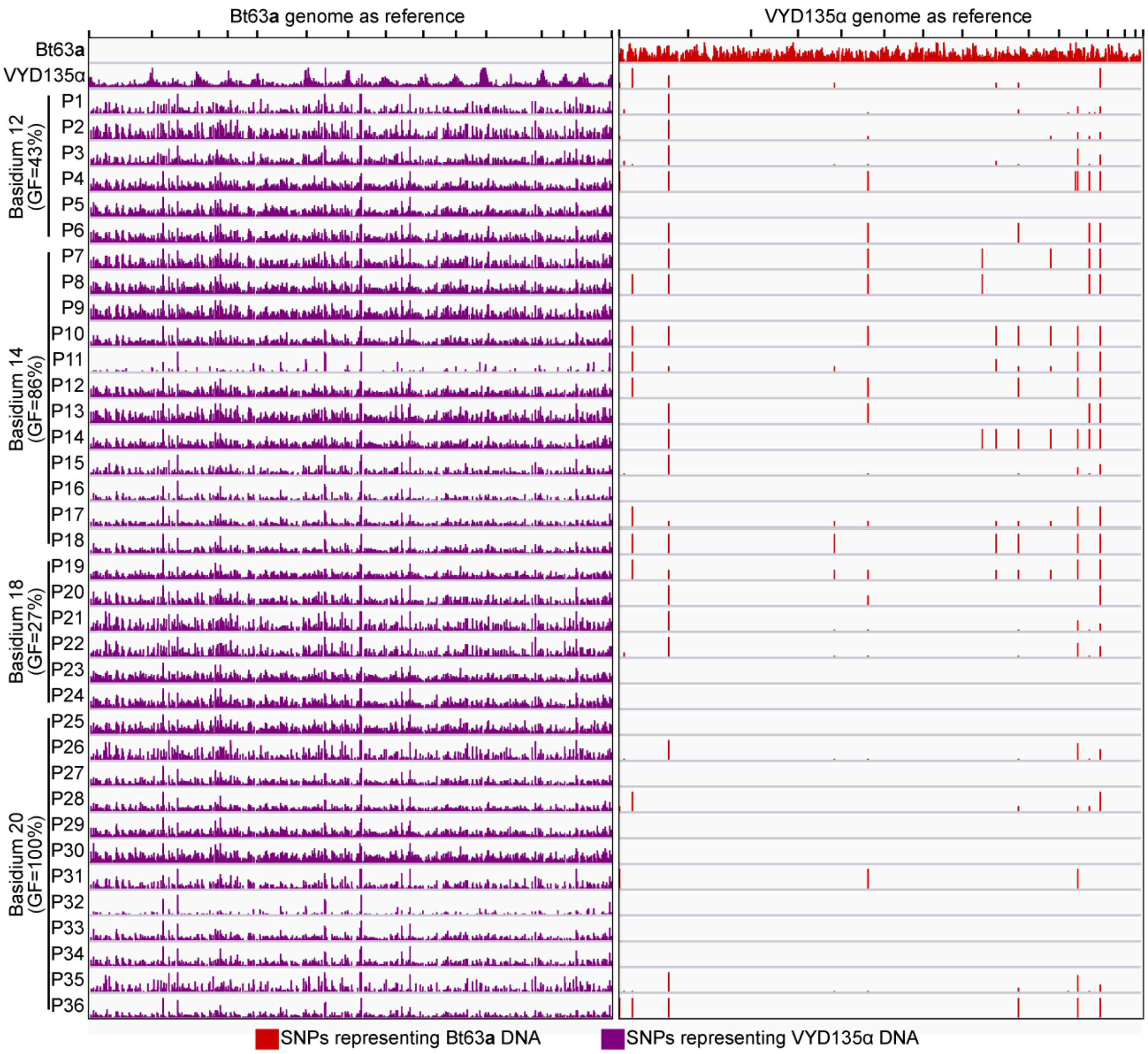


**Figure S5. VYD135α x Bt63a progeny lack signatures of meiotic recombination.** SNP analysis on VYD135α x Bt63**a** progeny revealed no contribution of the Bt63**a** parental genome in the progeny as evidenced by the presence of SNPs only against Bt63**a** (left panel) but not against VYD135α genome (right panel). The few SNPs observed in VYD135 as well as all VYD135α x Bt63**a** progeny lie within nucleotide repeat regions. GF stands for germination frequency and P stands for progeny.

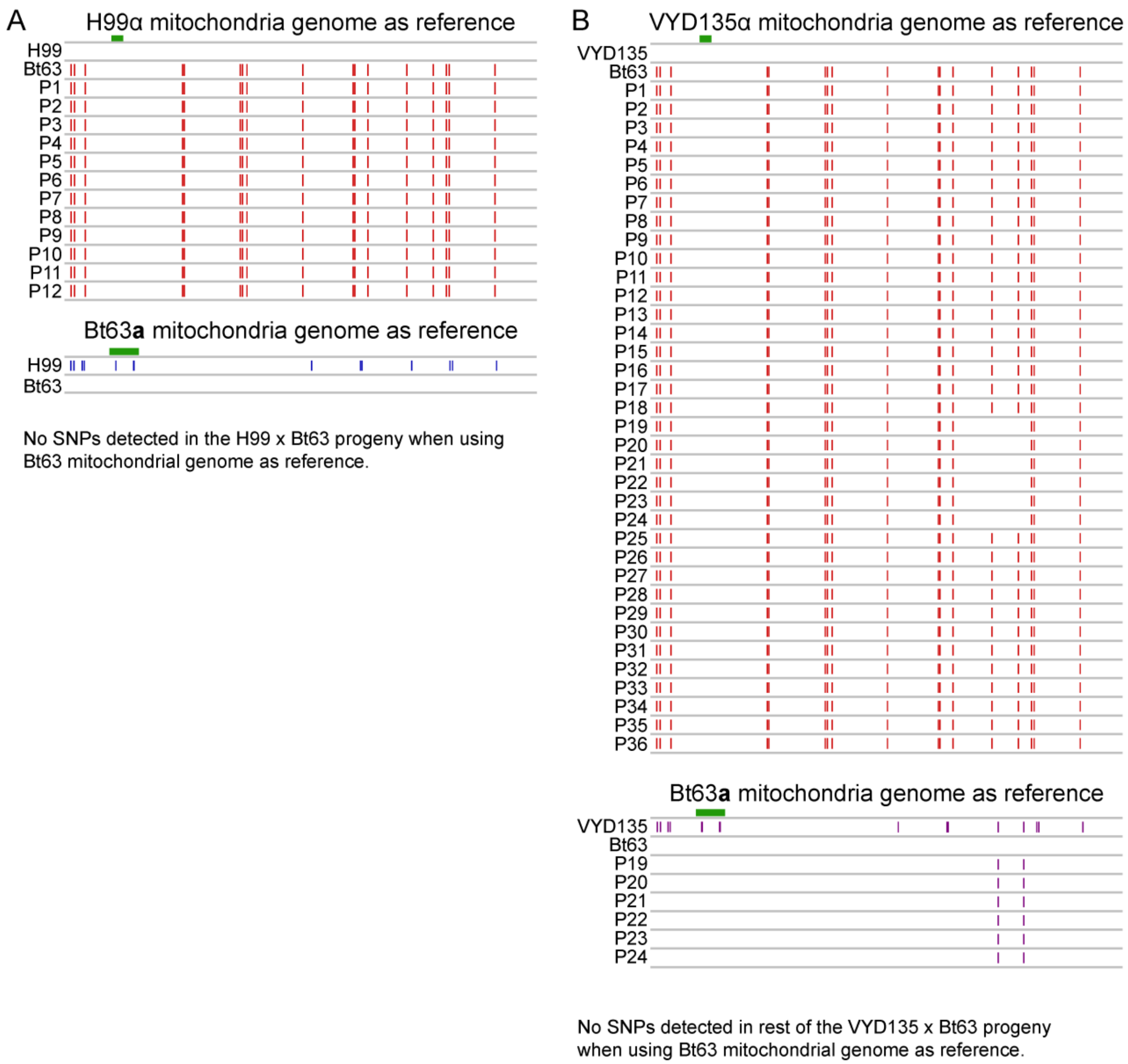


**Figure S6. Mitochondria are inherited from *MAT*a parent in all of the progeny. (A)** A map of SNPs detected in H99α x Bt63**a** progeny when using H99α mitochondrial DNA (upper panel) and Bt63**a** mitochondrial DNA (lower panel) as the reference. **(B)** SNP analysis revealed variants in all the progeny when using VYD135α mitochondrial DNA as the reference but not when using Bt63**a** mitochondrial DNA. The two SNPs detected against Bt63**a** DNA in progeny P19-24 (Basidium 18) suggest recombination of two parental mitochondrial DNA during mating. The green bar in each panel depicts the fragment used for PCR analysis in figure S3. P stands for progeny.

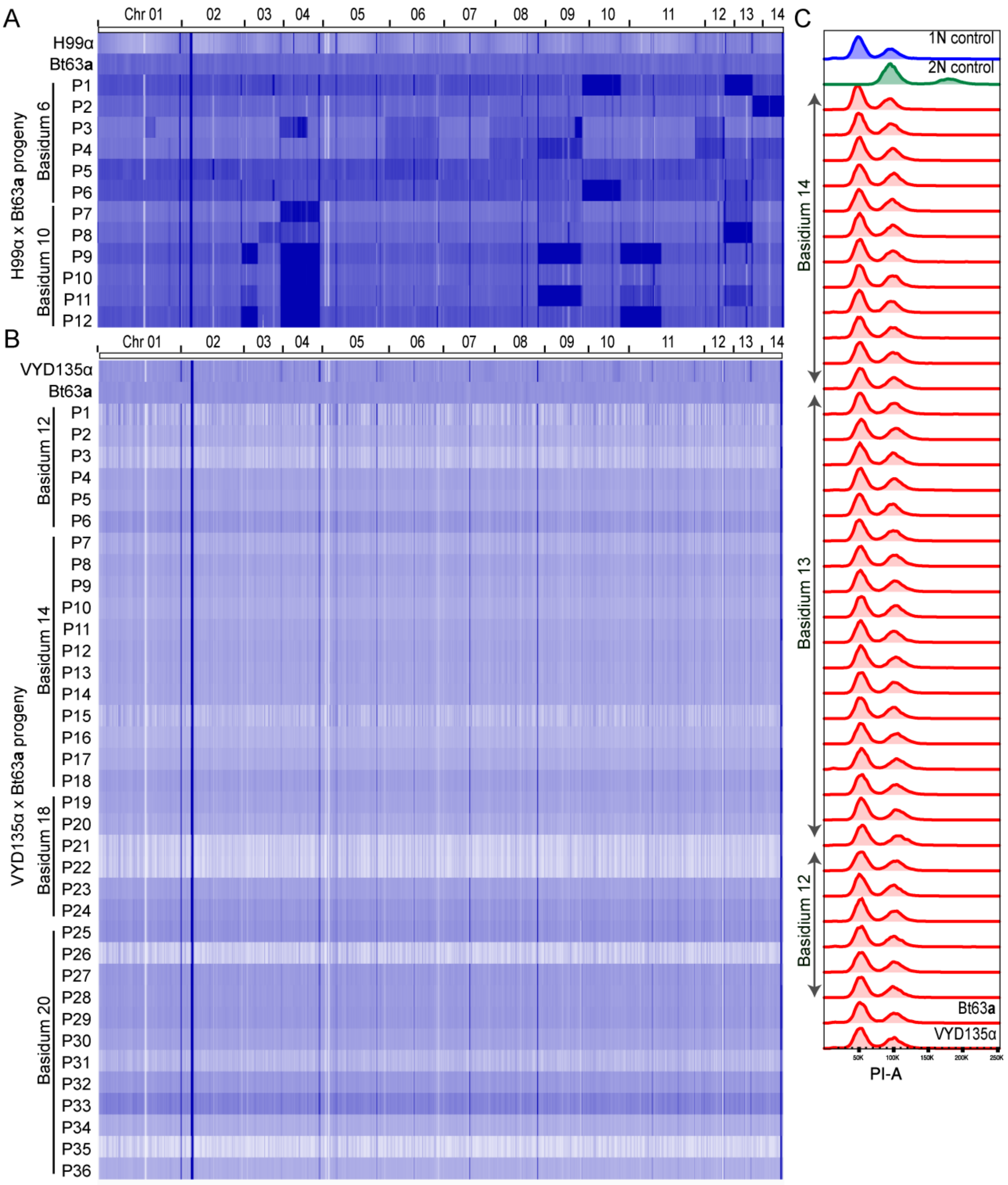


**Figure S7. VYD135α x Bt63a progeny are haploid. (A)** Whole-genome sequencing of the H99α x Bt63**a** progeny revealed extensive aneuploidy in the progeny. Each progeny seemed to exhibit aneuploidy for at least one chromosome. **(B)** Whole-genome sequencing data revealed that the progeny obtained from VYD135α x Bt63**a** 5-week old crosses are euploid in nature as they show a uniform level of genomic content when mapped to the Bt63 genome. VYD135α and Bt63**a** whole-genome sequencing data were also mapped as controls. Each lane represents one strain, and the difference in intensity correlates with the number of reads obtained per sample. **(C)** Flow-cytometry analysis on progeny obtained from three basidia confirmed that all the germinating progeny are haploid. While progeny from B12 and B14 are the same as used for the whole-genome sequencing, progeny from B3 were subjected to only flow cytometry analysis. Bt63**a** and VYD135α were also analyzed as controls for this experiment. P stands for progeny.

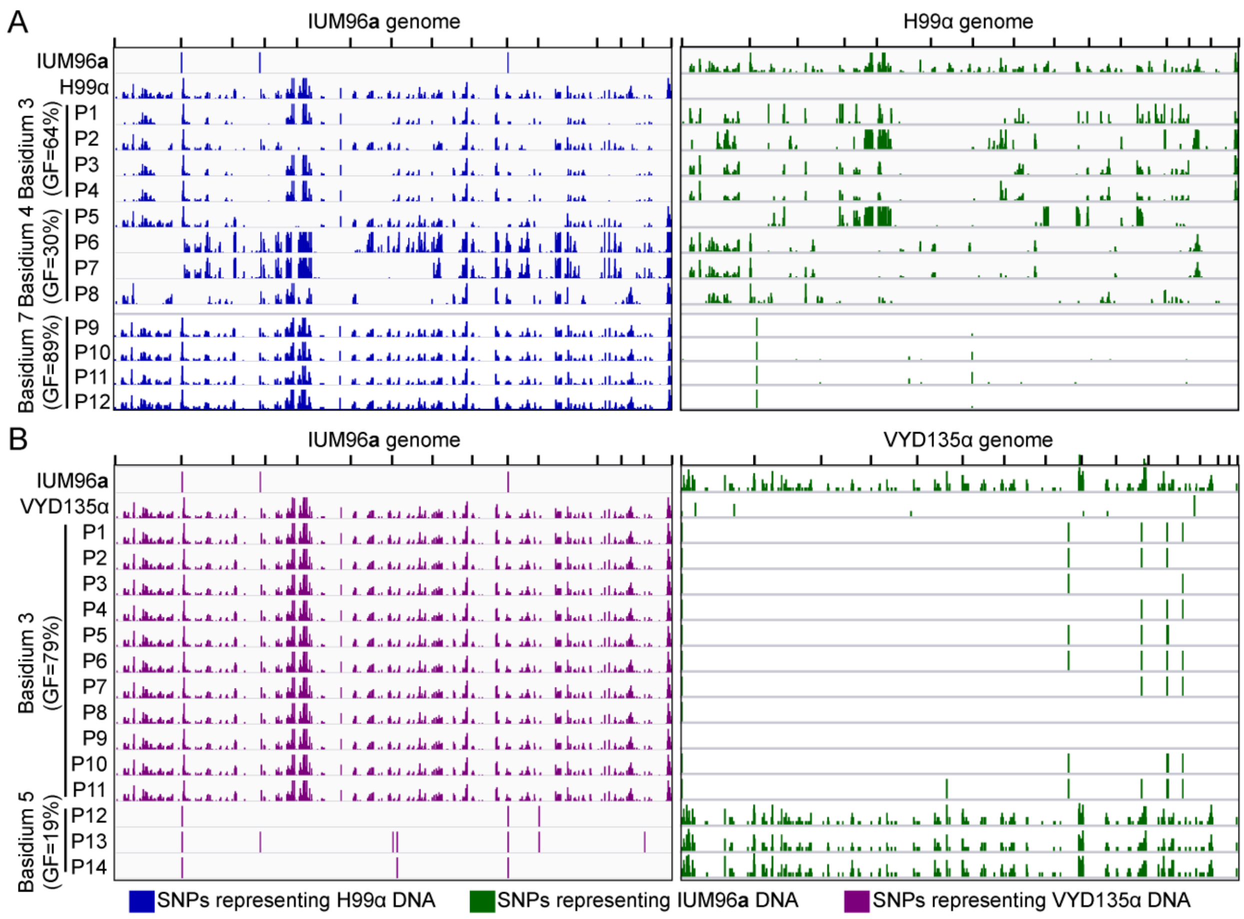


**Figure S8. IUM96a exhibits meiotic recombination in progeny with H99α but not with the genome shuffle strain VYD135α. (A)** The left panel depicts SNPs with respect to the IUM96**a** genome whereas the right panel shows SNPs against the H99α genome. Whole-genome sequencing, followed by SNP analysis, for the H99α x IUM96**a** progeny (basidia 3 and 4) revealed evidence of meiotic recombination in the progeny. Basidium 7 from the H99α x IUM96**a** cross produced uniparental progeny, which was confirmed by SNP analysis on a subset of these progeny. The progeny exhibited SNPs only against the IUM96**a** genome but not against the H99α genome. **(B)** SNP analysis from two different basidia revealed inheritance of only one set of parental nuclear DNA in the progeny from VYD135α x IUM96**a** cross. Basidium 3 progeny possessed DNA from only the VYD135α parent, while basidium 5 progeny inherited nuclear DNA from IUM96**a** alone. The results obtained from this analysis are congruent with mating-type PCR results shown in Table S2. GF stands for germination frequency and P stands for progeny.

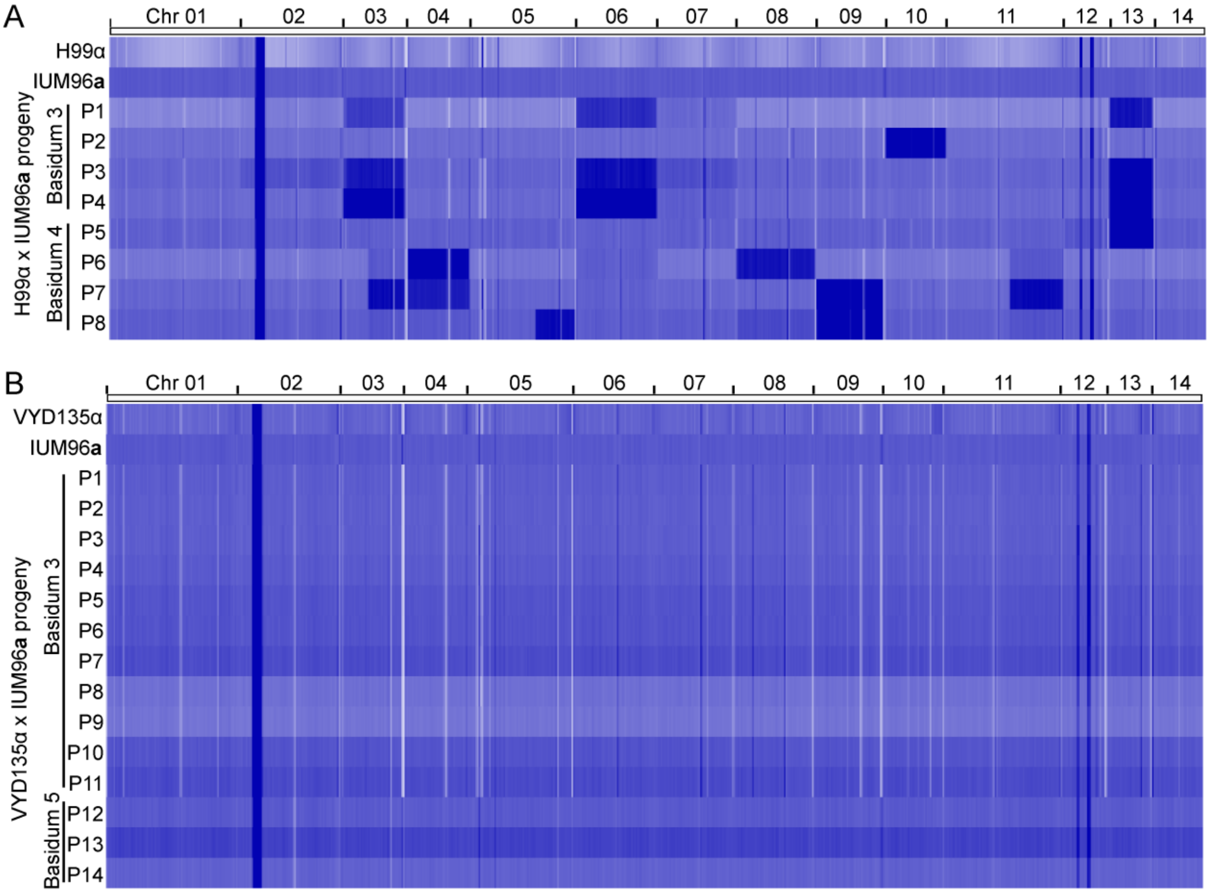


**Figure S9. Ploidy analysis of IUM96a progeny reveals haploid uniparental progeny.** Whole-genome sequencing analysis revealed the presence of multiple aneuploidies in the **(A)** H99α x IUM96**a** progeny, but a completely euploid genome for the **(B)** VYD135α x IUM96**a** progeny. P stands for progeny.

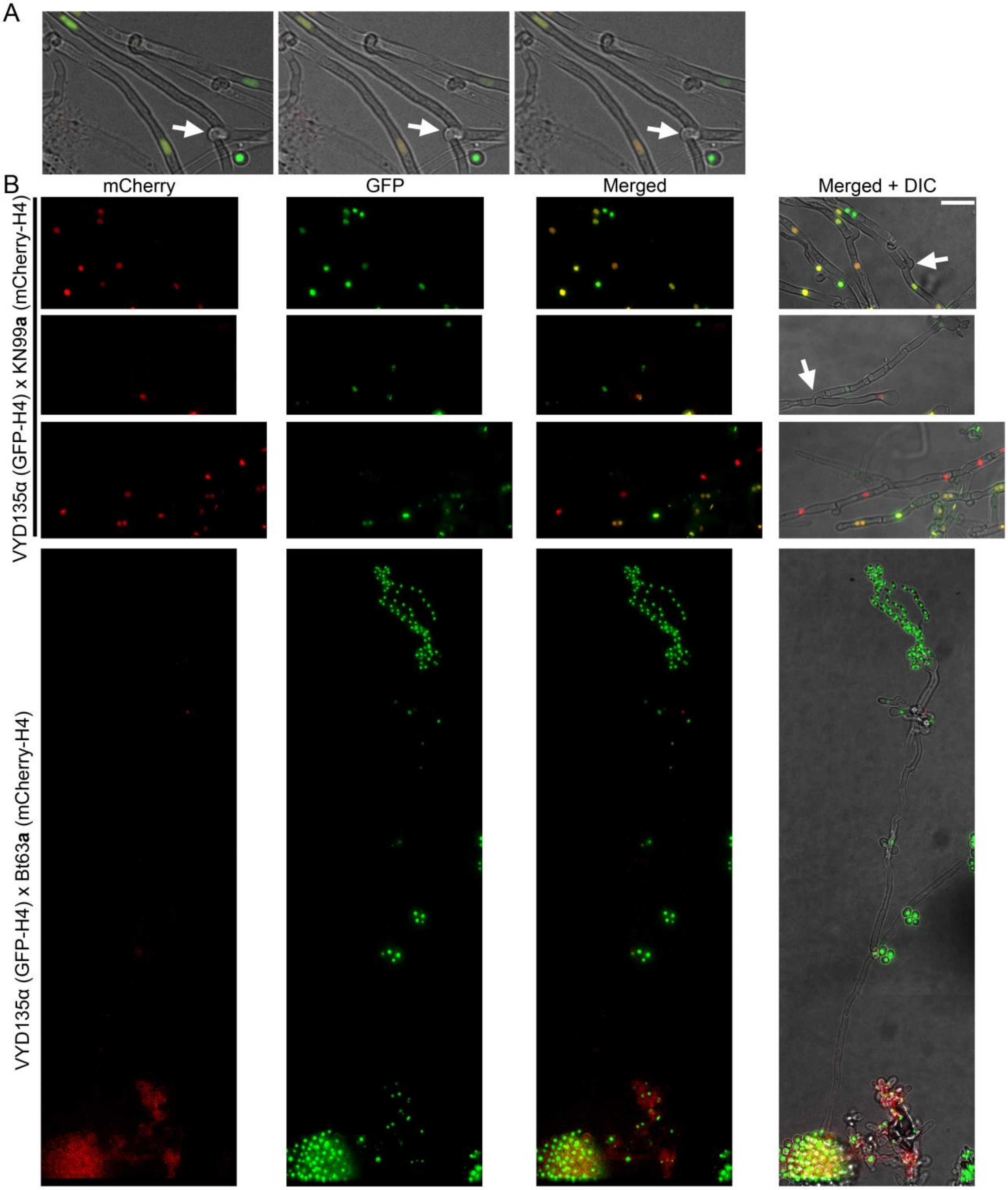


**Figure S10. Hyphal branches act as a gateway for nuclear separation during pseudosexual reproduction. (A)** Individual z-sections showing the hyphal branching (marked by arrow) where the two parental nuclei segregate in the figure 4B. **(B)** Images showing hyphal branching points where one of the parental nuclei separates from the main hyphae with two parental nuclei (Top two panels). The branch point is marked with the arrow. The lower two panels show the long hyphae with only one of the parental nuclei in them. The third panel shows other hyphae with both parental nuclei suggesting that separation occurred at an early stage. The fourth panel exhibits the same between VYD135α x Bt63**a** but also has a sporulating basidium on it. Bar, 10 µm.

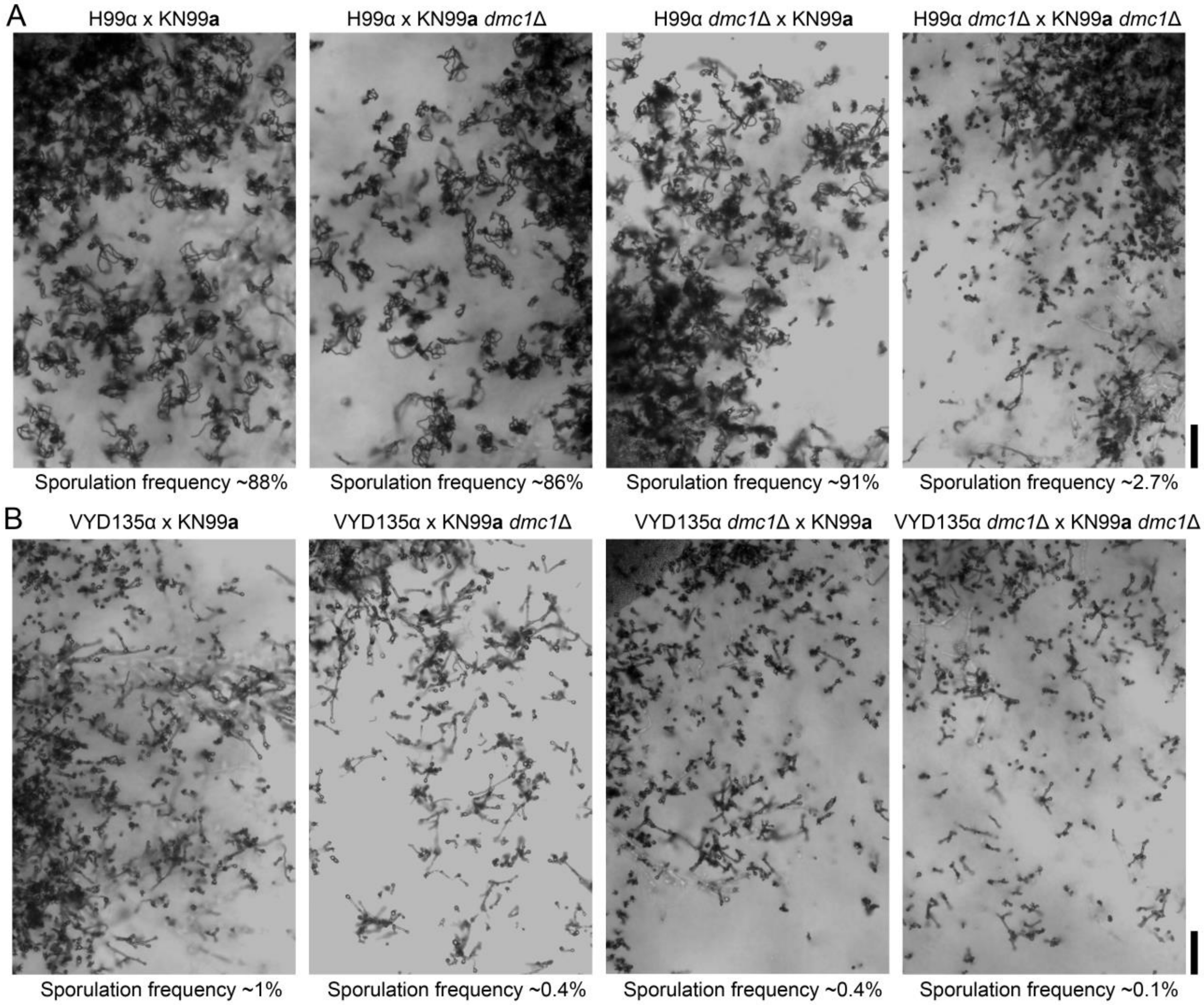


**Figure S11. Dmc1 deletion leads to severe sporulation defects in both sexual and pseudosexual reproduction.** Light microscopy images showing the phenotype of *DMC1* deletion in **(A)** H99α x KN99**a** unilateral crosses as well as bilateral mutant crosses and **(B)** VYD135α x KN99**a** *dmc1*Δ unilateral and bilateral crosses. The deletion of *DMC1* led to a reduction in sporulating basidia in bilateral mutant crosses.

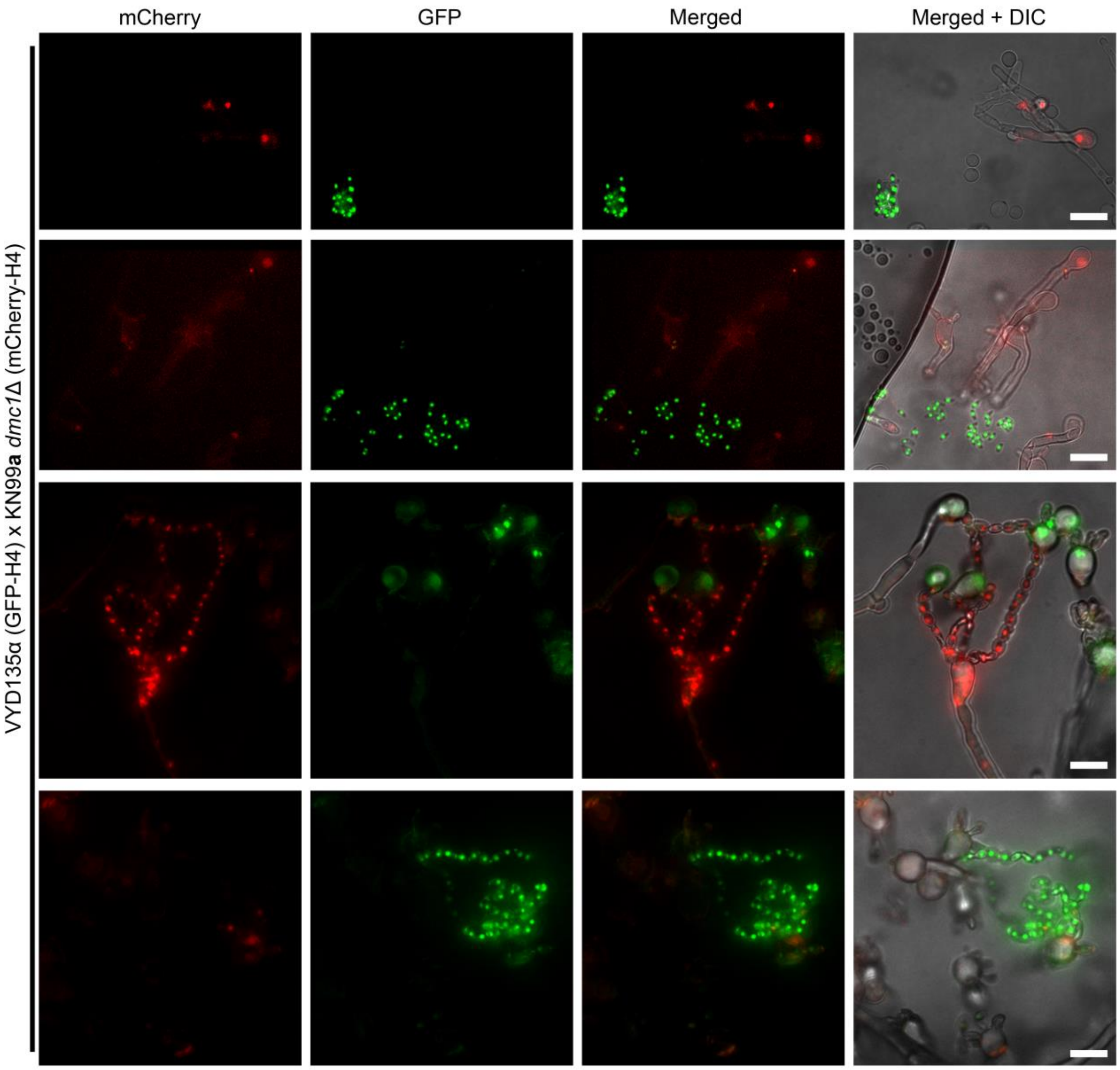


**Figure S12. Meiotic regulator Dmc1 is required for pseudosexual reproduction.** A cross between a GFP-H4 tagged VYD135α strain and an mCherry-H4 tagged KN99**a** *dmc1*Δ mutant revealed that Dmc1 is required for pseudosexual reproduction events. The majority of the KN99**a** *dmc1*Δ nucleus-containing basidia failed to produce spore chains (top two rows and bottom rows). While all 11 observed basidia with VYD135α nuclei produced spores, only 2 out of 19 observed basidia with KN99**a** *dmc1*Δ nuclei produced spores. One of these two is represented in the third row. Bars, 10 μm.
